# Apolipoprotein M attenuates anthracycline cardiotoxicity and lysosomal injury

**DOI:** 10.1101/2021.01.12.426397

**Authors:** Zhen Guo, Carla Valenzuela Ripoll, Antonino Picataggi, David R. Rawnsley, Mualla Ozcan, Julio A. Chirinos, Ezhilarasi Chendamarai, Amanda Girardi, Terrence Riehl, Hosannah Evie, Ahmed Diab, Attila Kovacs, Krzysztof Hyrc, Xiucui Ma, Aarti Asnani, Swapnil V. Shewale, Marielle Scherrer-Crosbie, Lauren Ashley Cowart, John S. Parks, Lei Zhao, David Gordon, Francisco Ramirez-Valle, Kenneth B. Margulies, Thomas P. Cappola, Ankit A. Desai, Lauren N. Pederson, Carmen Bergom, Nathan O. Stitziel, Michael P. Rettig, John F. DiPersio, Stefan Hajny, Christina Christoffersen, Abhinav Diwan, Ali Javaheri

**Affiliations:** Washington University School of Medicine, St. Louis, MO 63110; Perelman School of Medicine, University of Pennsylvania School of Medicine/Hospital of the University of Pennsylvania, Philadelphia, PA 19104; Hope Center, Washington University School of Medicine, St. Louis, MO 63110; John Cochran VA Medical Center, St. Louis, MO 63106; Beth Israel Deaconess, Harvard Medical School, Boston, MA 02115; Virginia Commonwealth University, Richmond, VA 23298 and Hunter Holmes McGuire VA Medical Center, Richmond, VA 23249; Wake Forest School of Medicine, Winston-Salem, NC 27104; Bristol Myers Squibb, Princeton, NJ 08543; Indiana University, Indianapolis, IN 46032; Dept. of Clinical Biochemistry, Rigshospitalet and Dept. of Biomedical Sciences, University of Copenhagen, 2100 Copenhagen, Denmark

**Keywords:** apolipoprotein M, anthracycline, autophagy, TFEB, cardiomyopathy

## Abstract

**Objectives:** Determine the role of apolipoprotein M (ApoM) in anthracycline (Dox) cardiotoxicity.

**Background:** ApoM binds the cardioprotective sphingolipid sphingosine-1-phosphate (S1P). Circulating ApoM is inversely associated with mortality in human heart failure (HF).

**Methods:** In the Penn HF Study (PHFS), we tested the relationship between ApoM and mortality in a subset with anthracycline-induced cardiomyopathy. We measured ApoM in humans and mice treated with Dox and utilized hepatic ApoM transgenic (*Apom^TG^*), ApoM knockout (*Apom^KO^*), ApoM knock-in mice with impaired S1P binding, and S1P receptor 3 (S1PR3) knockout mice in Dox cardiotoxicity. We assayed autophagy in left ventricular tissue from anthracycline-induced HF patients versus donor controls.

**Results:** ApoM was inversely associated with mortality in PHFS, and Dox reduced circulating ApoM in mice and breast cancer patients. *Apom^TG^* mice were protected from Dox-induced cardiac dysfunction and loss of left ventricular mass. *Apom^TG^* attenuated Dox-induced impairment in autophagic flux in vivo and accumulation of insoluble p62, which was also observed in the myocardium of patients with anthracycline-induced HF. In vehicle-treated mice, ApoM negatively regulated transcription factor EB (TFEB), a master regulator of autophagy and lysosomal biogenesis. The effect of ApoM on TFEB required both S1P binding and S1PR3. In the presence of Dox, ApoM preserved TFEB and cardiomyocyte lysosomal abundance assessed as lysosomal associated membrane protein 1 positive structures in vivo, while S1P mimetic pretreatment of cardiomyocytes prevented Dox-induced changes in lysosomal pH.

**Conclusions:** ApoM attenuates Dox cardiotoxicity via the autophagy-lysosome pathway. The association between ApoM and reduced mortality may be explained by its role in sustaining autophagy.

**Highlights:**

- Circulating ApoM is inversely associated with survival in human anthracycline-induced cardiomyopathy
- Anthracycline treatment reduces circulating ApoM in humans and mice
- Increasing ApoM attenuates doxorubicin cardiotoxicity, lysosomal injury and preserves myocardial autophagic flux, but does not impact doxorubicin anti-neoplastic efficacy
- Autophagic impairment is characteristic of human anthracycline cardiomyopathy

## Introduction

Chemotherapeutic treatments improve survival for many patients with cancer, but these improvements are offset by increases in therapy-related cardiovascular mortality (1). Anthracyclines, such as doxorubicin (Dox), are utilized for the treatment of cancers including breast cancer, leukemia, lymphoma, and sarcoma; however, the use of anthracyclines is limited by acute and chronic cardiotoxicity. The use of Dox remains common, particularly in children, with nearly 50% of children with cancer treated with an anthracycline (2). In children with acute lymphoblastic leukemia treated with at least one dose of Dox, 57% have cardiovascular abnormalities in follow-up (3). Although cardiotoxicity is clinically defined as a reduction in left ventricular ejection fraction (LVEF), reduced LV mass is a clinical sign that portends a poor prognosis (4), independent of changes in LVEF or body weight (5). In Long-term, anthracycline cardiotoxicity can result in heart failure (HF) in approximately 2% of patients, a diagnosis that carries a grave prognosis (6–8). Proposed mechanisms of Dox cardiotoxicity include DNA damage downstream of topoisomerase 2b (9), mitochondrial iron overload (10) and lysosomal injury with decreased nuclear translocation of transcription factor EB (TFEB), a master regulator of autophagy and lysosomal biogenesis (11,12).

High-density lipoprotein (HDL) has been suggested as a potential therapeutic for Dox cardiotoxicity and HF (13,14). In vitro studies indicate that the HDL attenuates cell death due to Dox via sphingosine-1-phosphate (S1P), a bioactive sphingolipid that binds G protein-coupled receptors on cardiomyocytes (15). Over 70% of S1P binds directly to the lipocalin apolipoprotein M (ApoM) on HDL particles, while the remainder of S1P is associated with albumin (16). ApoM is secreted primarily by hepatocytes, and it is associated with ~5% of HDL particles (and < 2% of low-density lipoprotein particles) (17–19).

ApoM exerts pleiotropic effects, including antioxidant and anti-atherogenic actions (20,21), regulation of inflammation (21), endothelial protection (16,22), and increased cell survival (23). Hepatocyte-specific overexpression of human *APOM* in mice (*Apom^TG^*) results in significant increases in plasma ApoM and S1P (19), while ApoM knockout mice (*Apom^KO^*) exhibit 50% reductions in plasma S1P, with normal amounts of S1P on albumin (16).

Our prior clinical observations demonstrate reduced circulating ApoM is associated with increased mortality in HF patients (24). In addition, we recently observed that reduced ApoM is closely linked to adverse effect of diabetes on human HF with preserved ejection fraction (25,26). In HF patients, ApoM is inversely associated with mortality independent of HF subtype, ischemic heart disease, HDL-C, or apolipoprotein A-I (ApoA-I), the main protein constituent of HDL (24). Despite these observations, little is known about the in vivo mechanisms by which ApoM may improve HF survival. In the present study, we hypothesized that increasing ApoM would attenuate Dox cardiotoxicity. Using anthracycline cardiotoxicity as a model, we identify that ApoM is a novel regulator of the autophagy-lysosome pathway. These studies point towards a novel function of the ApoM/S1P axis in regulating the autophagy-lysosome pathway in the myocardium and could explain the observed associations between ApoM and HF survival.

## Methods

### Reagents

Doxorubicin hydrochloride (50 mg) was purchased from United States Pharmacopeia (USP, R11760, Rockville, MD, USA) and dissolved in 20 mL of molecular grade water to get a 2.5 mg/mL stock, which was used in animal and cell experiments.

### Rodent studies

All murine studies were approved by the Institutional Animal Care and Usage Committee at the Washington University in St. Louis or by the Danish Animal Experiments Inspectorate and were performed following the Guide for the Care and Use of Laboratory Animals. Mice were maintained on a 12:12-hr light-dark schedule in a temperature controlled specific pathogen-free facility and fed standard laboratory mouse chow. *Apom^TG^* and *Apom^KO^* mice were previously described (16,19). *Myh6-Cre* mice were purchased from the Jackson Laboratory (Bar Harbor, ME, USA) (27), while *S1pr3^KO^* mice were previously described (28). Detailed methods for generation of Rosa-LAMP1-RFP mice and ApoM-knockin control or triple mutant mice (ApoM-CTR or ApoM-TM, respectively) are provided in the supplement. In age and sex-matched adult mice, cardiotoxicity was induced by intraperitoneal (IP) injection of Dox at specified doses. Acute models utilized 20 mg/kg Dox one dose, while chronic models utilized 5 mg/kg once weekly x 4 weeks as previously described (11,12). Single dose IP injections were also performed with 10 mg/kg. Echocardiography was performed as previously described (29), and read blindly by a cardiologist. LV mass and EF were obtained using the Vevostrain package in Vevo (Fujifilm Visualsonics, Toronto, Canada). Leukemic mice were generated by IV injection of male mice with 5 x 10^5^ APL cells (30). After allowing 4 days for engraftment, animals were treated with Dox on days 4, 6, 8, 11, 13, and 15 at 4 mg/kg subcutaneously. On day 17, peripheral blood was obtained via the retroorbital plexus under anesthesia and white blood cell (WBC) counts obtained using an automated cell counter (Hemavet 950, Drew Scientific). The percentage of circulating APL cells in blood was determined by flow cytometry.

### Histologic analyses

Histological analyses were performed as previously described (29). Electron microscopy was performed as previously described (31).

### Cytoplasm and Nuclear protein extraction

Cardiac tissues were fractionated into nucleus-enriched and cytoplasmic samples by using a CelLytic™ NuCLEAR Extraction kit (Sigma, Nxtract), as previously described (32).

### Insoluble protein isolation

Insoluble protein isolation was performed as previously described (29).

### Quantitative Real-Time Polymerase Chain Reaction analysis (qPCR)

Quantitative Real time PCR was performed as described (29).

### Western blot analysis

Western blot was performed on mice tissue or human samples as previously described (33).

### Human samples and ApoM determination

Circulating ApoM protein determination was performed from two independent cohorts. First, patients > 18 years of age diagnosed with HER2-positive breast cancer and scheduled to receive adjuvant therapy (anthracyclines, taxanes, and trastuzumab) were recruited prospectively and consecutively from Massachusetts General Hospital, MD Anderson Cancer Center, and McGill University (34,35). The study was approved by the institutional review board of the participating institutions, and all subjects provided informed consent. The typical cancer treatment regimen consisted of Dox 60 mg/m^2^ and cyclophosphamide 600 mg/m^2^ every 3 weeks for 4 cycles. At 3 months, all patients received paclitaxel 80 mg/m^2^ and trastuzumab 2 mg/kg weekly for 12 weeks, followed by trastuzumab 6 mg/kg every 3 weeks for 1 year. ApoM was measured in plasma samples using the SomaScan aptamer-based proteomics platform, as previously described and validated (24,36,37). Second, to determine the relationship between ApoM protein levels and mortality in anthracycline cardiotoxicity, we included patients from the Penn Heart Failure Study (PHFS) who had a history of anthracycline-induced cardiomyopathy. ApoM determination in this cohort was previously described (24).

For assessment of insoluble p62 in the human myocardium, de-identified human heart tissue samples were obtained from the Human Heart Tissue Bank at the University of Pennsylvania as previously described (38). Transmural LV samples from non-failing donors (obtained from brain-dead donors with no clinical history of HF) or anthracycline-induced cardiomyopathy (obtained at time of orthotopic heart transplantation) were obtained from the LV free wall, with epicardial fat excluded, and flash frozen in liquid nitrogen.

### Statistical analyses

All data are presented as Mean ± SEM or median ± 95% CI and analyzed by GraphPad Prism 9.0. The statistical comparisons between two groups were analyzed by unpaired Student’s t-test. The statistical significance among multiple groups was analyzed by analysis of variance (ANOVA) and multiple testing corrections as specified in each figure legend. In all of the analyses, a value of *P*□<□0.05 was considered statistically significant. Cox proportional hazard models were used to assess the univariate relationship between ApoM and survival in PHFS.

### Study approval

All animal studies were approved by the Animal Studies Committee at Washington University School of Medicine. Human studies were deemed exempt by the Washington University School of Medicine Institutional Review Board due to the fact that only de-identified human samples were utilized.

Additional methodology is described in the online supplement.

## Results

### ApoM is inversely associated with mortality in anthracycline cardiomyopathy

We previously reported that circulating ApoM is inversely associated with survival in HF patients (24). Whether ApoM is associated with mortality in patients with anthracycline cardiomyopathy is presently unknown. To test this, we identified patients in the PHFS with a history of anthracycline cardiomyopathy (n=46). Patient characteristics are shown in **Supplementary Table 1**. Among patients with a history of anthracycline cardiomyopathy, 11 patients died during follow-up. Baseline ApoM levels below the median were associated with reduced survival (**Figure 1A**). In univariate Cox proportional hazard models, we observed an inverse association between baseline ApoM and risk of death (**Figure 1B**), standardized hazard ratio 0.49 (95% CI 0.25-0.88, p=0.02).

**Figure 1.**
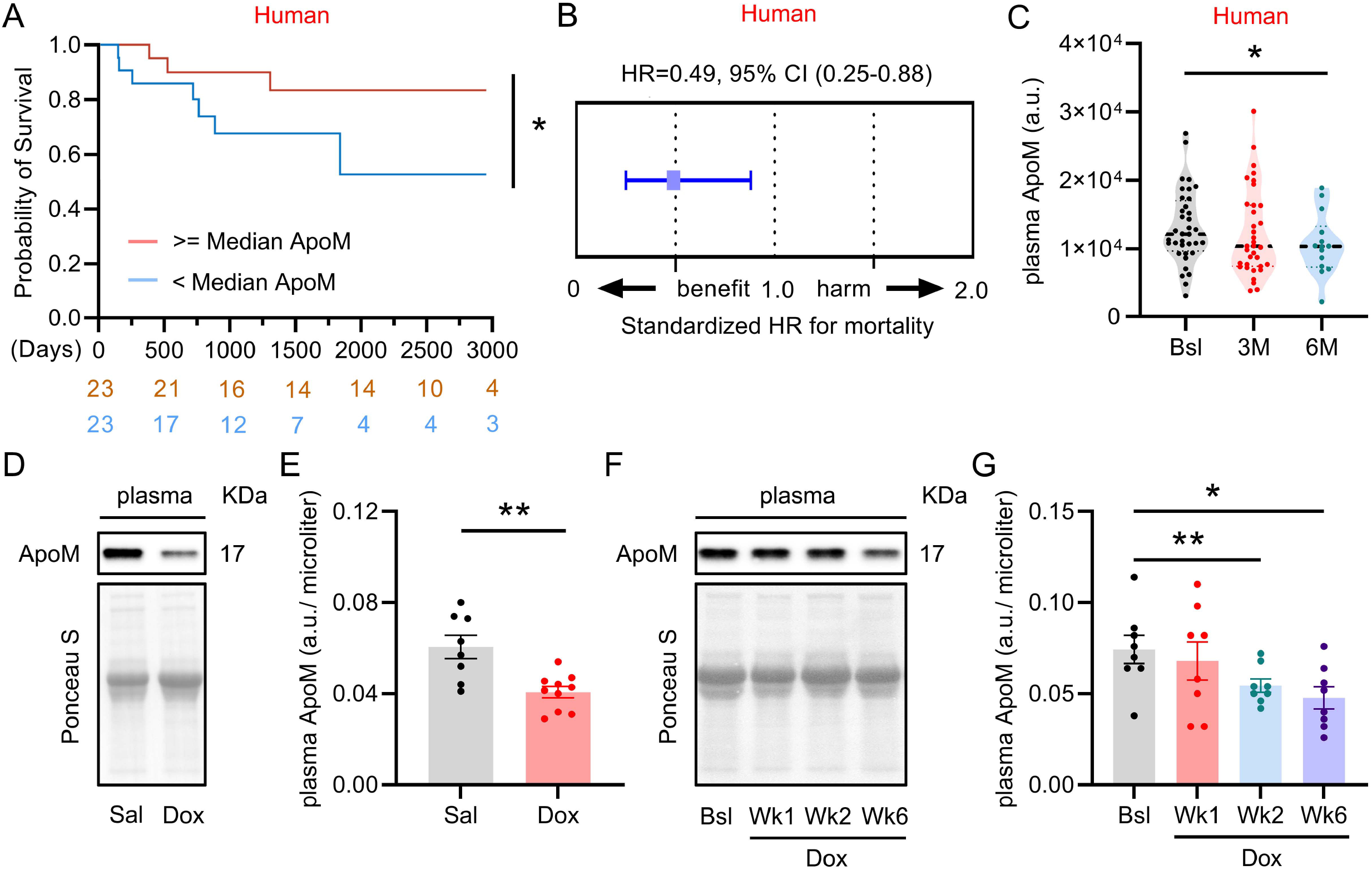
ApoM is inversely associated with mortality in anthracycline cardiomyopathy. **A)** Kaplan-Meier survival curves for all-cause mortality. The number of patients at risk at each time point is presented below the graph. **B)** Hazard ratio per standard deviation ApoM in patients with anthracycline cardiomyopathy. Cox proportional hazard model, n=46. **C)** Aptamer-based ApoM plasma levels in patients with breast cancer after 4 cycles of anthracycline. Mixed-effects model with Dunn’s correction for multiple comparisons, n=36 at baseline (Bsl), 35 at 3 months (3M) and 14 at 6 months (6M). Each dot represents one patient. Dotted lines represent median and quartiles. **D)** Representative western blot for ApoM from plasma isolated from mice 48 hr after treatment with saline (Sal) or 10 mg/kg doxorubicin (Dox) IP. **E)** Quantification of (D). Student’s t-test, n=8 Sal vs n=10 Dox. Each dot represents one mouse. **F)** Representative western blot for ApoM from plasma isolated from mice after treatment with Sal or 5 mg/kg Dox IP. Wk1 means 48 hr after 1^st^ dose, Wk2 means 48 hr after 2^nd^ dose and Wk6 means 6 weeks after 1^st^ dose. **G)** Quantification of (F). One-way repeated measures ANOVA with Dunn’s correction for multiple comparisons, n=8 per group. Each dot represents one mouse. *p < 0.05, **p < 0.01.

Given the known cardiotoxicity caused by anthracyclines (39), we tested whether anthracyclines reduce ApoM in humans and mice. We observed decrease in circulating ApoM in breast cancer patients (clinical information shown in **Supplementary Table 2**) treated with anthracyclines for four cycles (**Figure 1C**). In murine studies, we observed parallel reductions in circulating ApoM after acute Dox administration (**Figure 1D-E**), an effect that was dose and time-dependent (**Figure 1F-G**).

### ApoM attenuates doxorubicin induced cardiotoxicity

Given the reduction in plasma ApoM caused by Dox, we investigated whether increasing ApoM was sufficient to attenuate Dox cardiotoxicity. In initial studies comparing *Apom^TG^* to non-transgenic littermate controls, we modeled cardiotoxicity using high dose, acute treatment with Dox (20 mg/kg IP, **Figure 2A**). In these experiments, we observed that *Apom^TG^* mice were protected from Dox-induced reductions in LV mass and heart weight/tibia length (HW/TL) that occurred in littermate controls (**Figure 2B-C**). In chronic studies with Dox delivered via tail vein injection (5 mg/kg x 4 doses delivered once per week, **Figure 2D**), we observed reductions in LVEF in littermate controls but not in *Apom^TG^* mice (**Figure 2E**). In chronic models, littermate control mice, but not *Apom^TG^*, demonstrated significantly reduced HW/TL (**Figure 2F**).

**Figure 2.**
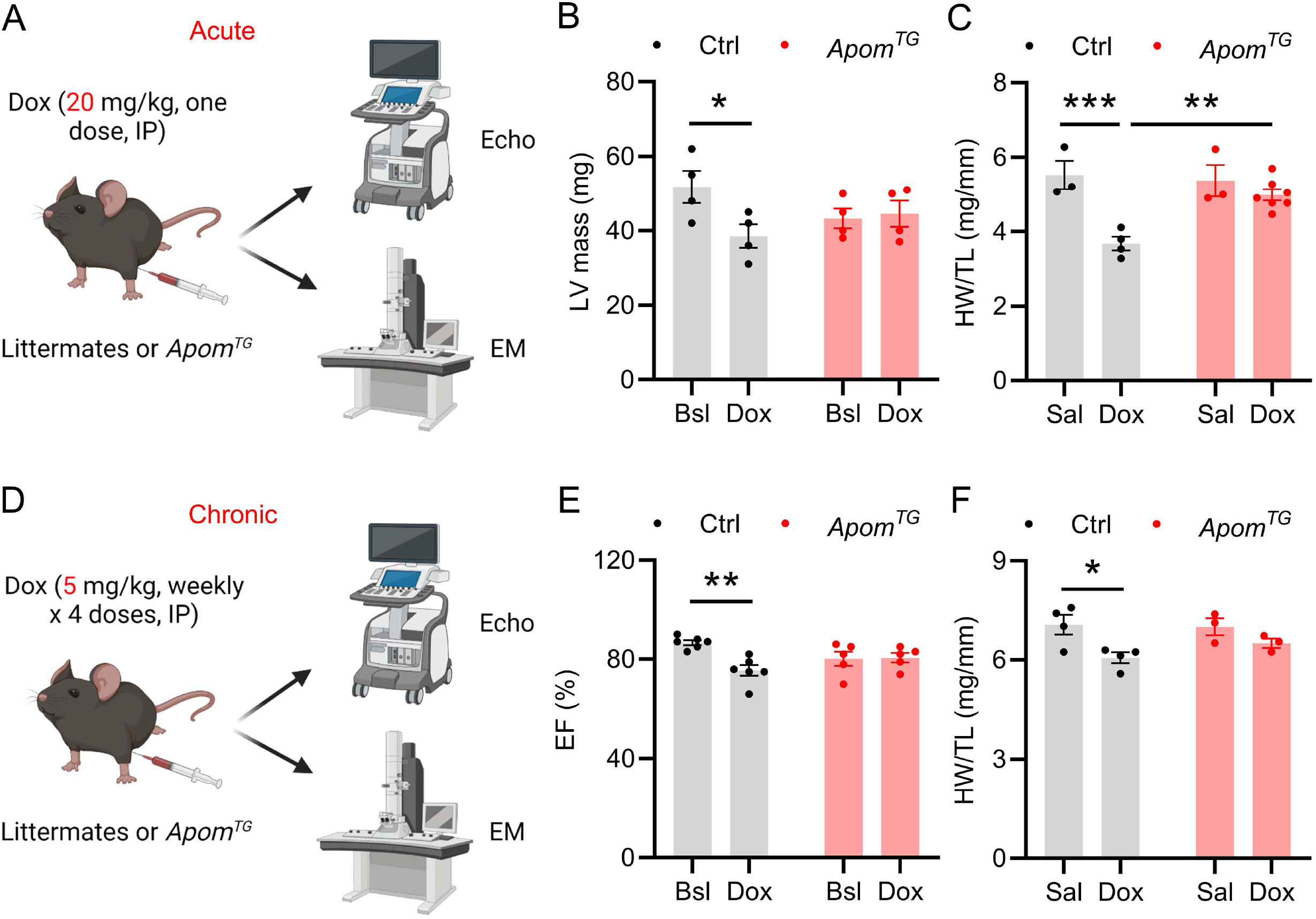
ApoM attenuates doxorubicin induced mortality and cardiotoxicity. **A)** Schematic diagram of the acute experimental model. **B)** Change in LV mass from baseline (Bsl) and 5 days after 20 mg/kg doxorubicin (Dox) IP. Two-way repeated measures ANOVA with Sidak’s correction for multiple comparisons, n=4 per group. **C)** Change in heart weigh/tibia length (HW/TL) from saline (Sal) and 5 days after 20 mg/kg Dox IP. Two-way ANOVA with Sidak’s correction for multiple comparisons, n=3~7. **D)** Schematic diagram of the chronic experimental model. **E)** Change in LVEF from Bsl and 3 months after 4 weekly doses of 5 mg/kg Dox IV. Two-way repeated measures ANOVA with Sidak’s correction for multiple comparisons, n=6 Ctrl vs n=5 *Apom^TG^.* **F)** Change in HW/TL from Sal and 3 months after 4 weekly doses of 5 mg/kg Dox IV. Two-way ANOVA with Sidak’s correction for multiple comparisons, n=3~4. Each dot represents one mouse. *p < 0.05, **p < 0.01, ***p < 0.001.

Finally, using a murine model of acute promyelocytic leukemia (30,40), we observed that Dox treatment (subcutaneous 4 mg/kg x 6 doses, **Supplementary Figure 1A**) extended survival to a similar degree in *Apom^TG^* mice compared to littermate controls (**Supplementary Figure 1B**) with blast suppression 2 days after the last dose of Dox (**Supplementary Figure 1C**), indicating that ApoM does not attenuate the anti-leukemic efficacy of Dox in vivo.

### ApoM does not alter DNA damage, apoptosis, fibrosis, or Akt signaling due to doxorubicin

We next investigated mechanisms by which ApoM could attenuate Dox cardiotoxicity. In these studies, we utilized an intermediate dose of Dox (10 mg/kg) sufficient to result in signaling changes without significant weight loss. We examined myocardial γ-H2AX (41) foci as a marker of DNA damage resulting from acute Dox administration (48 hr after one dose, 10 mg/kg), and found no difference in foci in *Apom^TG^* versus littermate control mice (**Supplementary Figure 2**). As HDL can attenuate apoptosis caused by Dox in vitro (15), we performed anti-caspase 3 immunohistochemistry in myocardial sections from the above mice, and again found no significant differences in active, nuclear caspase 3 (42) (**Supplementary Figure 3**).

Furthermore, the levels of myocardial anti-caspase 3 staining were relatively modest compared to positive controls obtained from prior closed-chest ischemia reperfusion experiments (43). Evaluation of mitochondrial ultrastructure by electron microscopy did not demonstrate marked qualitative differences in acute or chronic Dox cardiotoxicity models between *Apom^TG^* and littermate controls (**Supplementary Figure 4A-B**), nor was there evidence of reduced fibrosis by Masson’s trichrome stain in the chronic model (**Supplementary Figure 5**).

Transgenic overexpression of ApoA-I (*Apoa1^TG^*), the main protein constituent of HDL, attenuates Dox cardiotoxicity via the phosphoinositol-3-kinase/protein kinase B (PI3K/Akt) pathway (14). We confirmed that *Apoa1^TG^* mice exhibited increased Akt phosphorylation at Ser473; however, we did not observe increased phosphorylation of Akt at Ser473 in *Apom^TG^* mice (**Supplementary Figure 6A-D**). Altogether, these studies suggest that ApoM did not attenuate cardiotoxicity in Dox cardiotoxicity by impacting DNA damage, apoptosis, Akt signaling, or fibrosis in the myocardium.

### ApoM overexpression prevents Dox-induced autophagic impairment

We therefore turned our attention to whether ApoM might be affecting autophagy and lysosome dysfunction, which have recently been reported in murine models of Dox cardiotoxicity (11,12). To measure autophagic flux, we performed immunohistochemistry for autophagy proteins LC3, a marker of autophagosomes, and p62, an autophagy receptor and substrate, respectively, from myocardial sections of mice treated with vehicle and Dox plus or minus the lysosomal inhibitor chloroquine. Compared to littermate controls, *Apom^TG^* exhibited intact autophagic flux after Dox, as evidenced by accumulation of LC3 (**Figure 3A-B**) and p62 (**Figure 3C-D**) with chloroquine.

**Figure 3.**
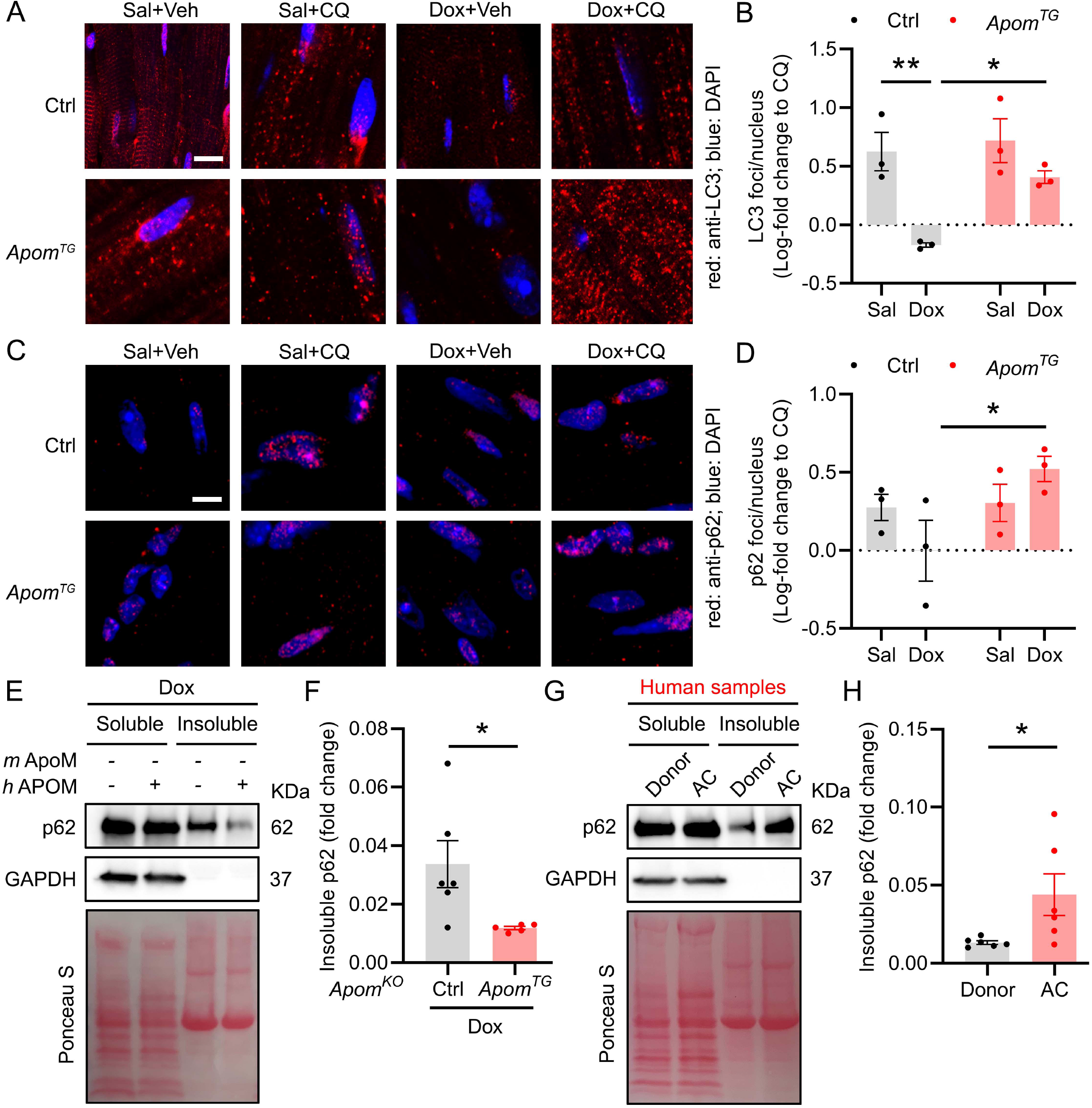
ApoM attenuates doxorubicin-induced autophagic impairment. **A)** Representative images of immunohistochemical staining of mid-myocardial sections with α-LC3 and DAPI in littermate control (Ctrl) and *Apom^TG^* mice 48 hr post-treatment with saline (Sal) or doxorubicin (Dox, 10 mg/kg) IP and, 4 hr prior to euthanasia, with vehicle (Veh) vs chloroquine (CQ, 60 mg/kg) IP, scale bar=5 μm. **B)** Blinded quantification of LC3 foci from (A). Two-way ANOVA with Sidak’s correction for multiple comparisons, n=3 per group. Data were log transformed due to violation of normality. **C)** Representative images immunohistochemical staining of mid-myocardial sections with α-p62 and DAPI in mice from (A), scale bar=5 μm. **D)** Blinded quantification of p62 foci from (C). Two-way ANOVA with Sidak’s correction for multiple comparisons, n=3 per group. Data were log transformed due to violation of normality. **E)** Representative western blot for myocardial soluble vs insoluble p62 from *Apom^KO^* and *Apom^KO^ Apom^TG^* mice 5 days after 15 mg/kg Dox IP. **F)** Insoluble p62 quantified relative to soluble p62. Student’s t-test, n=6 *Apom^KO^* vs n=5 *Apom^KO^ Apom^TG^*. **G)** Representative western blot for myocardial soluble vs insoluble p62 from patients with a history of heart failure due to anthracycline cardiomyopathy (AC) versus donor controls (Donor) obtained from patients without any known clinical heart failure. **H)** Insoluble p62 quantified relative to soluble p62. Student’s t-test, n=6 per group. Each dot represents one mouse or one patient. *p < 0.05, **p < 0.01.

To verify the effect of ApoM on autophagy, we generated a new line of mice with human ApoM overexpression in the murine ApoM knockout background (*Apom^KO^Apom^TG^*) and assayed accumulation of insoluble p62, an indicator of autophagic impairment that occurs in human and murine HF (29,44). In contrast to Dox-treated *Apom^KO^* mice that accumulated insoluble p62, *Apom^KO^Apom^TG^* mice exhibited reduced insoluble p62 (**Figure 3E-F**). We then confirmed that accumulation of insoluble p62 is a pathophysiological feature of human anthracycline cardiomyopathy, as left-ventricular myocardium from anthracycline cardiomyopathy patients exhibited increased insoluble p62 relative to donors without a history of HF (**Figure 3G-H**).

### ApoM regulates transcription factor EB in an S1P-dependent manner

Reductions in autophagy due to Dox may be causally related to the decrease in active, nuclear TFEB, a transcription factor and master regulator of lysosomal biogenesis. To determine the effects of ApoM on nuclear translocation of TFEB, we performed nuclear protein isolation from control and *Apom^TG^* hearts. Though ApoM overexpression did not alter total TFEB protein levels (**Supplementary Figure 7A-B**), *Apom^TG^* mice demonstrated unexpected reductions in active, nuclear TFEB at baseline (**Figure 4A-B**), as evidenced by reductions in the faster migrating, dephosphorylated band that represents active TFEB. Nevertheless, in response to Dox, *Apom^TG^* mice were resistant to further decreases in nuclear TFEB that occurred in littermate controls (interaction p<0.001, **Figure 4C-D**), suggesting that in the context of ApoM overexpression, Dox has a differential effect on nuclear TFEB content.

**Figure 4.**
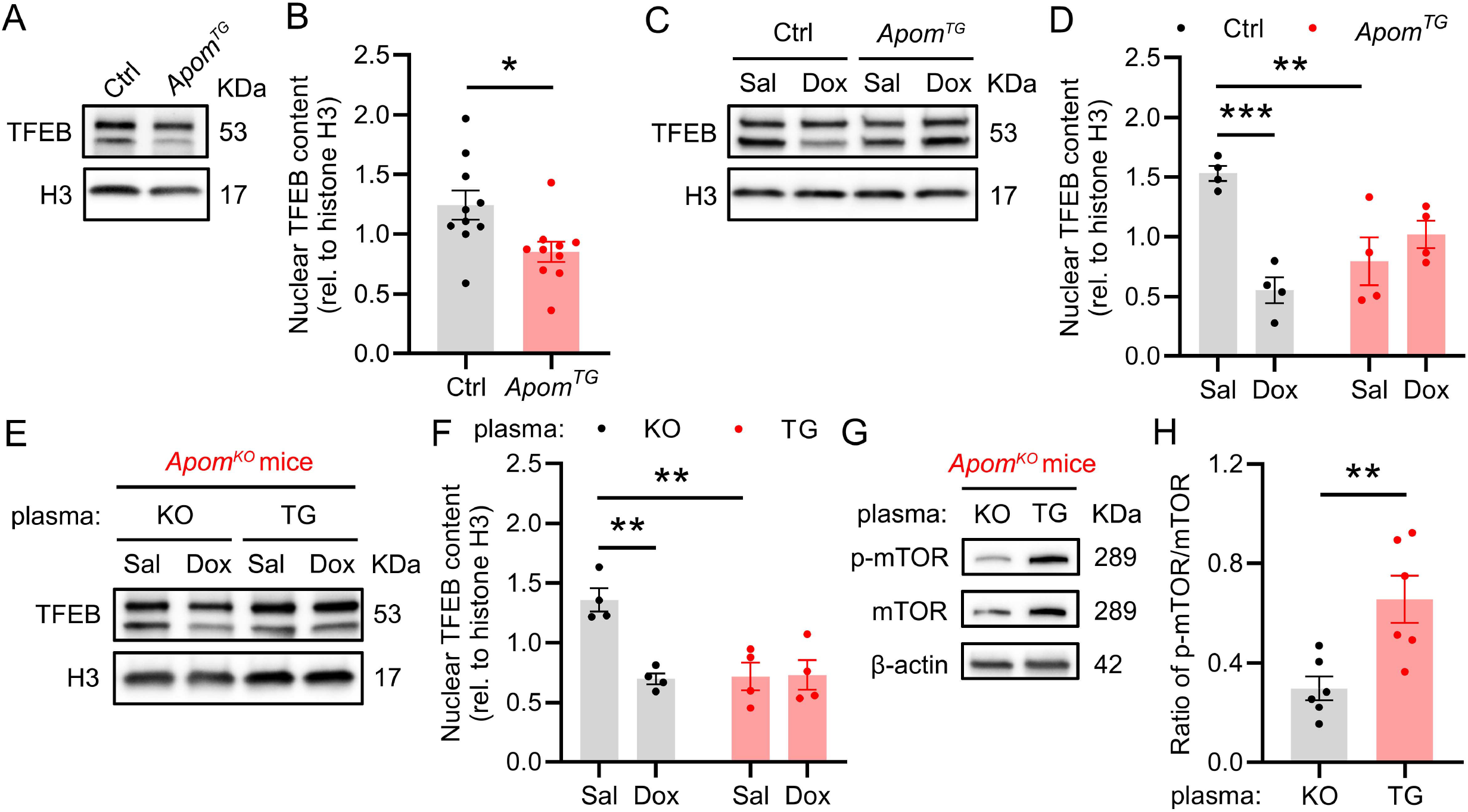
ApoM attenuates doxorubicin-induced reductions in nuclear TFEB. **A)** Representative western blot of TFEB from isolated myocardial nuclear protein extracts from littermate control (Ctrl) and *Apom^TG^* mice. **B)** Quantification of nuclear TFEB from (A). Student’s t-test, n=10 per group. **C)** Representative western blot of TFEB from isolated myocardial nuclear protein extracts from Ctrl and *Apom^TG^* mice 48 hr after treatment with saline (Sal) or doxorubicin (Dox, 10 mg/kg) IP. **D)** Quantification of nuclear TFEB from (C). Two-way ANOVA with Sidak’s correction for multiple comparisons, n=4 per group. **E)** Representative western blot of TFEB from isolated myocardial nuclear protein extracts of *Apom^KO^* mice 2 days after transfer of 120 μL plasma obtained from *Apom^KO^* or *Apom^TG^* donor mice 48 hr after treatment with Sal or Dox (10 mg/kg) IP. **F)** Quantification of nuclear TFEB from (E). Two-way ANOVA with Sidak’s correction for multiple comparisons, n=4 per group. **G)** Representative western blot of p-mTOR and mTOR from myocardial obtained from *Apom^KO^* mice 2 days after transfer of 120 μL plasma obtained from *Apom^KO^* (KO) or *Apom^TG^* (TG) donor mice. **H)** Quantification of p-mTOR/mTOR from (G). Student’s t-test, n=6 per group. Each dot represents one mouse. *p < 0.05, **p < 0.01, ***p < 0.001.

To test whether acute increase in ApoM also alter nuclear TFEB content, we utilized plasma transfer from donor *Apom^TG^* or *Apom^KO^* mice to recipient *Apom^KO^* mice prior to administration of Dox. Plasma transfer increased human ApoM levels in *Apom^KO^* mice (**Supplementary Figure 7C**). *Apom^TG^* plasma also reduced baseline TFEB levels (that were elevated in *Apom^KO^* mice receiving *Apom^KO^* plasma) and attenuated further decrease in TFEB levels after Dox (interaction p<0.01, **Figure 4E-F**), similar to what we observed in *Apom^TG^* mice. Mechanistically, administration of *Apom^TG^* plasma resulted in increased phosphorylation of mammalian target of rapamycin (mTOR), the main negative regulator of TFEB, in myocardial tissue lysates (**Figure 4G-H**).

We next sought to determine whether our observation regarding TFEB regulation by ApoM requires canonical S1P-dependent signaling. To do so, we utilized a novel ApoM knock-in mouse (R98A, W100A, and R116A) called ApoM triple mutant (ApoM-TM) that cannot bind S1P. ApoM-TM mice demonstrated increased nuclear TFEB (**Figure 5A-B**), which phenocopied *Apom^KO^* mice (**Figure 5C-D**). Finally, knockout of S1PR3 also abrogated the effects of ApoM on TFEB (**Figure 5E-F**), suggesting that ApoM negatively regulates TFEB through S1P-mediated S1PR3 signaling.

**Figure 5.**
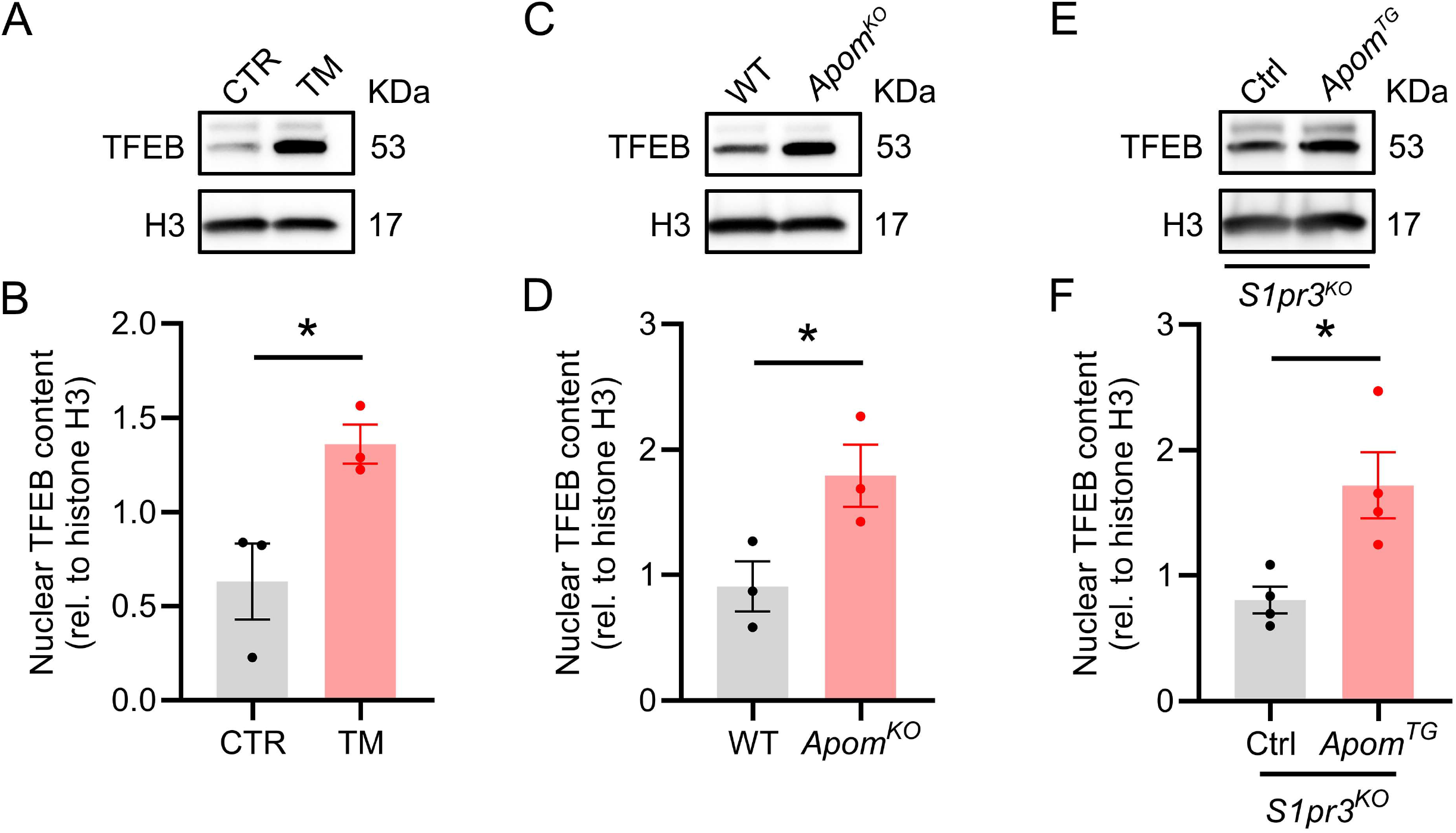
ApoM reduces nuclear TFEB in a S1P-dependent manner through S1PR3. **A)** Representative western blot of TFEB from isolated myocardial nuclear protein extracts from control (CTR) and ApoM triple mutant (TM) that cannot bind S1P. **B)** Quantification of nuclear TFEB from (A). Student’s t-test, n=3 per group. **C)** Representative western blot of TFEB from isolated myocardial nuclear protein extracts from wild type (WT) and *Apom^KO^* mice. **D)** Quantification of nuclear TFEB from (C). Student’s t-test, n=3 per group. **E)** Representative western blot of TFEB from isolated myocardial nuclear protein extracts in *Apom^TG^* mice crossed with *S1pr3^KO^* mice. **F)** Quantification of nuclear TFEB from (E). Student’s t-test, n=4 per group. Each dot represents one mouse. *p < 0.05.

### The role of ApoM and TFEB in anthracycline cardiotoxicity

To determine whether TFEB was causally involved in cardioprotection by ApoM, we utilized a validated strategy of adeno-associated virus 9 (AAV9) short-hairpin RNA-mediated TFEB knockdown (29). Littermate control and *Apom^TG^* mice were injected IV with 3.5e11 viral particles of AAV9-shScramble versus AAV9-shTFEB. AAV9-shTFEB transduction resulted in significant reductions in TFEB mRNA abundance in littermate controls and *Apom^TG^* mice (**Figure 6A**). TFEB knockdown had opposing effects on heart rate in littermate control vs *Apom^TG^* mice (interaction p<0.05, **Figure 6B**). While TFEB knockdown did not significantly affect ejection fraction (**Figure 6C**), *Apom^TG^* had significant increases in end-diastolic volume that did not occur in littermate controls (interaction p<0.05, **Figure 6D**); hence, in contrast to littermate controls, *Apom^TG^* mice suffered adverse LV remodeling after TFEB knockdown. To determine the effect of TFEB knockdown on autophagy, we performed immunostaining for LC3 in littermate control and *Apom^TG^* mice transduced with AAV9-shScramble or shTFEB. In comparison to mice transduced with shScramble, only *Apom^TG^* transduced with AAV9-shTFEB exhibited reductions in LC3 foci (**Figure 6E-F**). In littermate control wildtype mice, TFEB knockdown had no effect on LC3 foci, and TFEB knockdown was associated with increased heart rate, LV mass, the dV/dT max both in the presence or absence of Dox administration (p<0.05, **Figure 6G-I**). In contrast, when *Apom^TG^* AAV9-shTFEB mice were treated with Dox died after a single dose of Dox (5 mg/kg IP). In summary, *Apom^TG^* mice exhibit decreased nuclear TFEB, and are sensitive to further knockdown of TFEB, while knockdown of TFEB did not exacerbate and may even lessen echocardiographic markers of Dox cardiotoxicity.

**Figure 6.**
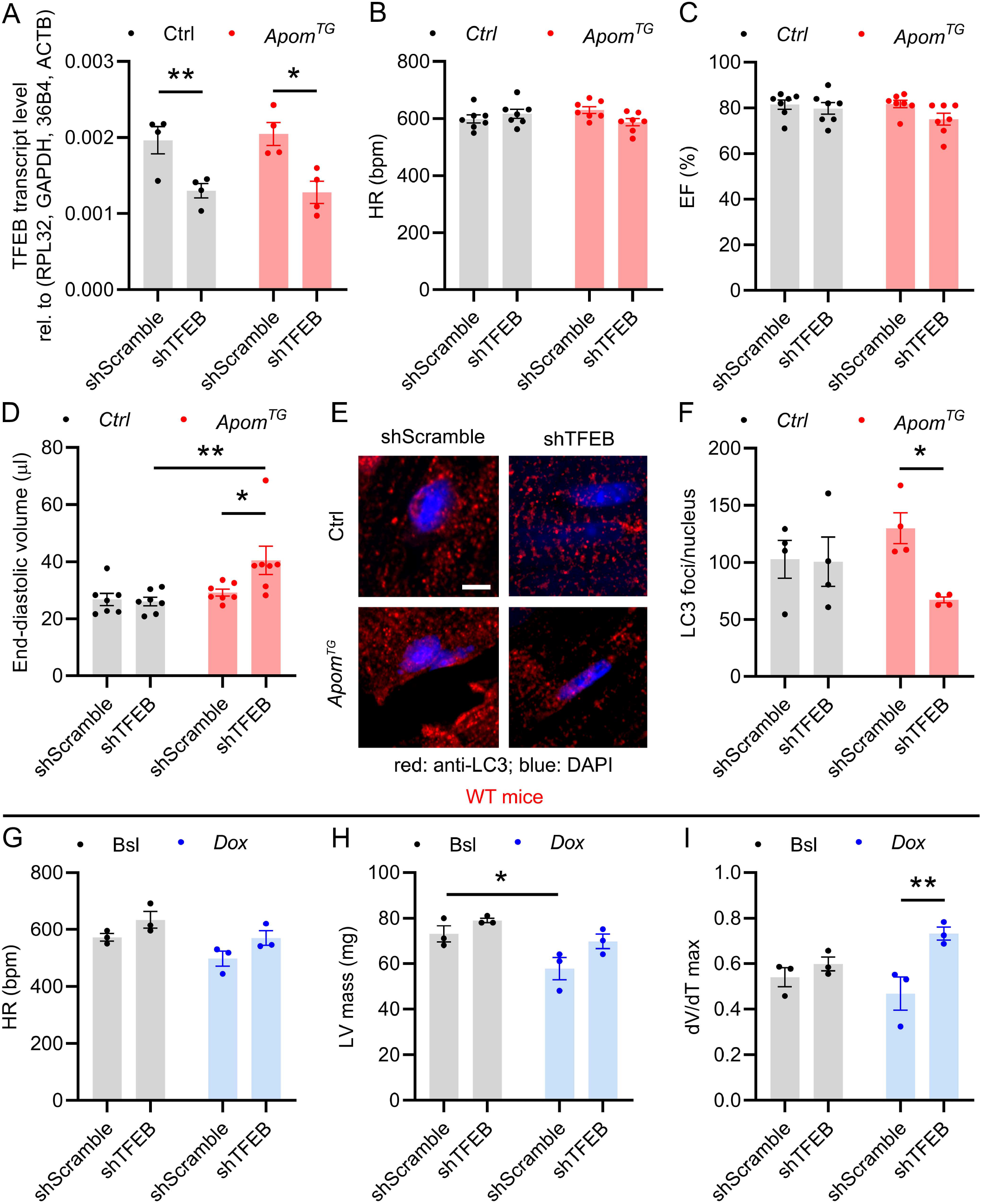
ApoM, TFEB, and anthracycline cardiotoxicity. **A)** TFEB mRNA abundance accessed by qPCR in littermate control (Ctrl) and *Apom^TG^* mice transduced with AAV9-shScramble or AAV9-shTFEB. Two-way ANOVA with Sidak’s correction for multiple comparisons, n=4 per group. **B-D)** Heart rate (HR), ejection fraction (EF), and end-diastolic volume assessed by echocardiography 2 weeks after viral transduction. Two-way ANOVA with Sidak’s correction for multiple comparisons, n=7 per group. **E)** Representative images of immunohistochemical assessment of LC3 foci in myocardial sections, scare bar=5 μm. **F)** Blinded quantification of LC3 foci from (E). Two-way ANOVA with Sidak’s correction for multiple comparisons, n=4 per group. **G-I)** HR, LV mass, and dV/dT assessed by echocardiography 2 weeks after AAV9-shScramble or AVV9-shTFEB viral transduction in wild type mice. Two-way repeated measures ANOVA with Sidak’s correction for multiple comparisons, n=3 per group. Each dot represents one mouse. *p < 0.05, **p < 0.01.

### ApoM and S1P attenuate Dox-induced lysosomal injury

Both anthracyclines and other chemotherapeutic drugs activate a lysosomal stress response characterized by loss of lysosomal pH (11) and lysosomal exocytosis, as evidenced by redistribution of lysosomes to the plasma membrane (45). Given the paradoxical findings that ApoM preserves autophagic flux after Dox but reduces baseline nuclear levels of TFEB, a transcription factor that regulates autophagy and lysosomal biogenesis, we reasoned that ApoM and S1P may protect against Dox-induced lysosomal injury in vivo. To test this, we utilized a lysosomal RFP reporter (*Myh6-Cre* LAMP1-RFP) to genetically label and track cardiomyocyte lysosomes (**Supplementary Figure 8A**). Expression of this LAMP1-RFP construct in cardiomyocytes does not significantly alter cardiac function or structure as measured by echocardiography (**Supplementary Figure 8B-C**), or significantly alter heart weight (**Supplementary Figure 8D**). We then treated *Myh6-Cre* Rosa-LAMP1-RFP mice with *Apom^TG^* plasma followed by Dox. Dox induced reductions in the RFP signal that were attenuated by *Apom^TG^* plasma (**Figure 7A-B**).

**Figure 7.**
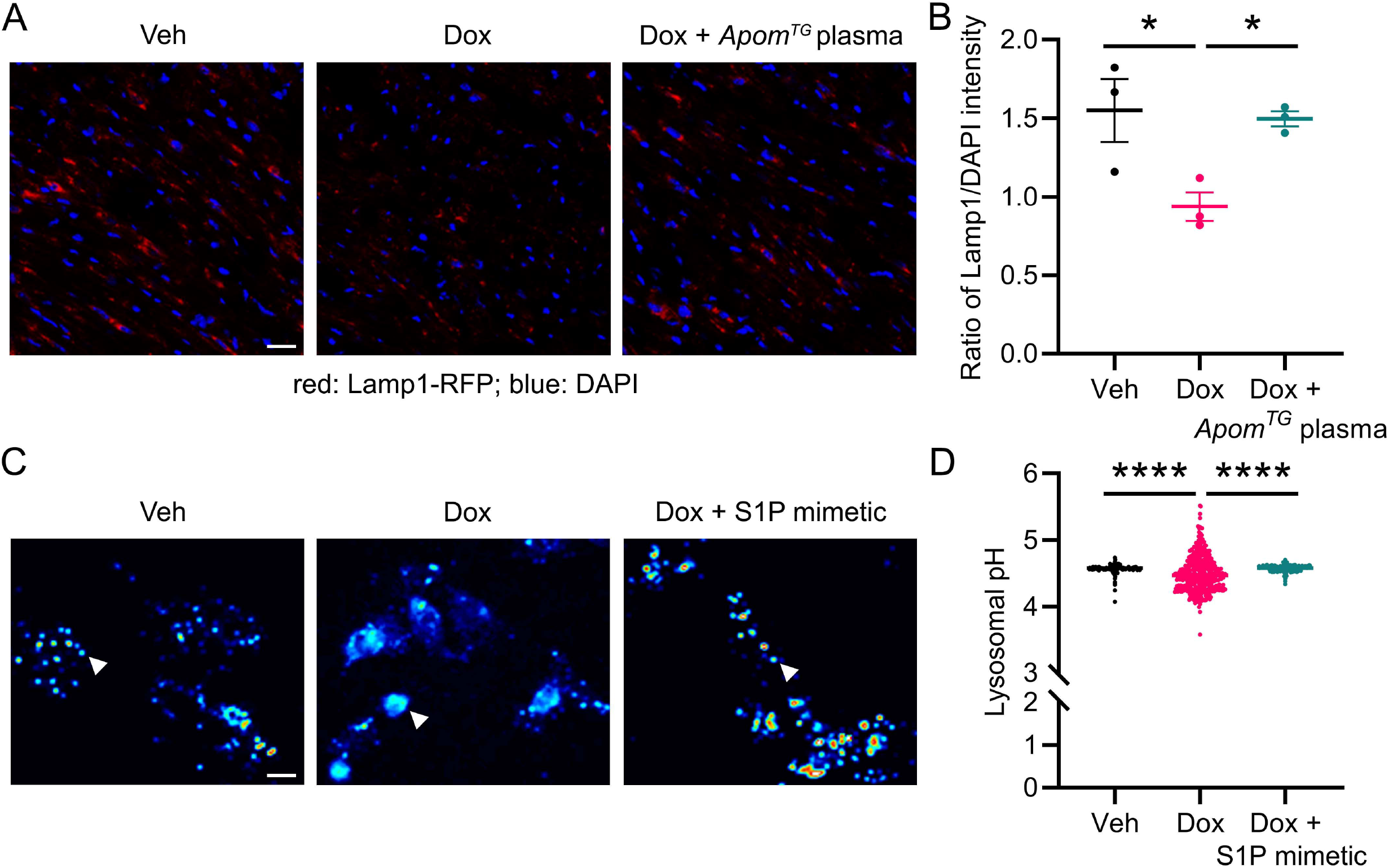
ApoM and S1P attenuate Dox-induced lysosomal injury. **A)** Representative images of red fluorescence of mid-myocardial sections in *Myh6-Cre* x Rosa-LAMP1-RFP mice injected with 120 μL plasma obtained from *Apom^TG^* mice, then treated with 2-days doxorubicin (Dox, 10 mg/kg) IP. **B)** Blinded quantification of red/blue intensity from (A). One-way ANOVA with Sidak’s correction for multiple comparisons. Each dot represents one mouse. **C)** S1P mimetic pre-treatment of neonatal rat cardiomyocytes attenuates lysosomal injury. Representative pseudocolor excitation ratio images of Lysosensor-loaded lysosomes in cells exposed to DMSO, 0.5 μM Dox and 0.5 μM Dox with 25 nM FTY720 pre-treatment for 5 min. Cells were incubated with 1 μM Lysosensor Yelow/Blue DND 160 for 3 min, washed with HCSS and imaged by collecting pairs of images excited at 380 nm and 340 nm through a long pass 480 nm emission filter through a 40 x /1.35 oil immersion lens (Olympus). Frequency distribution of the excitation ratio values in single lysosomes, scale bar=25 μm. White arrow indicates lysosome. **D)** Graph of lysosomal pH from (C) and data for individual lysosomes pooled from 3 separate litters of rats were collected in 30-60 individual cells. Kruskal-Wallis test with Dunn’s correction for multiple comparisons. Each dot represents one lysosome. *p < 0.05, ****p < 0.0001.

To further determine whether an S1P mimetic could attenuate lysosomal injury, we treated neonatal rat cardiomyocytes with either vehicle or the S1P mimetic FTY720 prior to treatment with Dox, followed by live cell microscopy with Lysosensor DND-160, a pH sensitive ratiometric probe that can be utilized to measure pH in acidified organelles. To harness only the S1P activating effects of FTY720, cardiomyocytes were pre-treated with 25 nM of FTY720 for 5 min prior to washing the cells with PBS and subsequent treatment with Dox. In these experiments, we were careful to treat with low-dose (0.5 μM) Dox for only 4 hr in order to minimize cytotoxic effects of the drug seen with higher doses and longer timepoints. Treatment of cardiomyocytes with Dox resulted in a characteristic pattern of lysosomal exocytosis that was abrogated by pre-treatment with FTY720 (**Figure 7C-D**). Furthermore, Dox resulted in acute changes in lysosomal pH over 4 hr that were attenuated by pre-treatment with FTY720. Hence, both ApoM and an S1P mimetic are able to protect cardiomyocyte lysosomes from Dox-induced injury and alterations in pH, respectively.

## Discussion

In this manuscript, we utilized a spectrum of human cohort and myocardial tissue samples as well as multiple murine models to demonstrate that ApoM attenuates the cardiotoxicity, autophagic impairment, and lysosomal injury that occurs due to anthracyclines. The translational relevance of our studies is highlighted by our observations that a) circulating ApoM is inversely associated with survival in human anthracycline-induced cardiomyopathy, b) anthracycline treatment reduces circulating ApoM in humans and mice, c) increasing ApoM attenuates Dox cardiotoxicity and preserves myocardial autophagic flux, but does not impact Dox anti-neoplastic efficacy, d) autophagic impairment is characteristic of human anthracycline cardiomyopathy. These findings are concordant with and expand upon our prior findings that ApoM is inversely associated with all-cause mortality in HF patients (24).

Mechanistically, our studies suggest ApoM, via S1P signaling, is a novel regulator of the autophagy-lysosome pathway and TFEB, both of which exhibit complex interactions in Dox cardiotoxicity. The main mechanistic conclusion of our work is that cardioprotection by ApoM and S1P (46,47) may be mediated both by TFEB, and by effects of ApoM/S1P on lysosomes and preservation of autophagic flux. Our findings further demonstrate that ApoM/S1P can protect lysosomes from Dox-induced injury. Across all our models, we consistently observe that untreated wildtype mice have higher nuclear TFEB levels than *Apom^TG^* mice; hence, it is likely that the effect of ApoM/S1P on lysosomes is not due to increased TFEB nuclear translation.

In fact, our data indicate two distinct effects of ApoM/S1P on TFEB. In the unstressed heart prior to Dox exposure, ApoM/S1P signaling negatively regulates TFEB nuclear translocation. Mutation of the ApoM-S1P binding pocket, ApoM knockout mice, and S1PR3 knockout all caused increases in myocardial nuclear TFEB content, suggesting that the ApoM/S1P axis is a rheostat for TFEB in the unstressed heart; i.e., in the absence of exogenous stress, increasing ApoM reduces active TFEB content, while reducing ApoM, or its ability to bind S1P or signaling through S1PR3, all increase nuclear TFEB. While our data point to the downstream events culminating in mTOR phosphorylation, further research will be needed to elucidate the precise mechanism by which ApoM/S1P/S1PR3 signaling can activate mTOR in cardiomyocytes. In contrast, in the setting of Dox, ApoM overexpression or delivery of plasma enriched for ApoM preserved both TFEB and lysosomal abundance in cardiomyocytes in vivo, as assayed by LAMP1-RFP positive structures. The effect on cardiomyocyte lysosomes was also observed in vitro after a brief incubation with an S1P mimetic. This relatively rapid effect of ApoM/S1P is also consistent with a post-transcriptional mechanism, although further research will be needed to identify precisely how ApoM/S1P can protect lysosomal structures.

The above studies along with our TFEB knockdown experiments suggest that a partial reduction in TFEB nuclear activity is well-tolerated and possibly protective, while further reduction of TFEB (at least in the context of ApoM overexpression) is likely toxic. Hence, there appears to be an optimal level of TFEB nuclear content: neither too low nor too high. Specifically, AAV9-shRNA mediated knockdown of TFEB only resulted in adverse cardiac consequences in *Apom^TG^*, but not littermate control mice. Further, TFEB knockdown in *Apom^TG^* mice decreased LC3 protein abundance, consistent with impaired autophagy. Importantly, the mRNA abundance of TFEB was reduced to an equal extent in both *Apom^TG^* and littermate control mice, consistent with post-transcriptional regulation of TFEB by ApoM/S1P signaling. Given the increased end-diastolic volume in *Apom^TG^* mice treated with AAV9-shTFEB in the absence of additional stress, it is unsurprising that these mice were intolerant of Dox. In contrast, TFEB knockdown in wildtype littermate control mice did not result in any enhanced sensitivity to Dox.

Another important finding of our studies is that autophagic impairment is a pathophysiological characteristic of anthracycline cardiomyopathy in humans. Although anthracyclines mediate toxicity via multiple pathways, lysosomes have recently emerged as a critical target of anthracyclines and other anti-neoplastic drugs (11,12,45), and have an increasingly recognized role in cardiometabolic disease (48). In humans with anthracycline cardiomyopathy, we observed accumulation of insoluble p62, suggesting that autophagic impairment observed previously in murine models (11) also occurs in humans. Our studies support that increasing ApoM might be an attractive cardioprotective strategy, as overexpression of ApoM did not impact the anti-neoplastic efficacy of Dox while protecting against cardiotoxicity and reducing insoluble levels of p62. In addition, since preservation of autophagic flux has been implicated in the pathogenesis of HF more broadly (29,44), our studies may help explain the broader link between ApoM and HF outcomes.

### Perspectives

What Is New?

- ApoM is inversely associated with survival in patients with anthracycline cardiomyopathy.
- Anthracycline treatment reduces circulating ApoM in humans.
- Increasing ApoM attenuates doxorubicin-induced cardiotoxicity and lysosomal injury.

What Are the Clinical Implications?

- Anthracycline-induced reductions in ApoM may be an important mediator of subsequent cardiotoxicity.
- ApoM increasing therapies may attenuate doxorubicin cardiotoxicity.
- Further studies must assess whether increasing ApoM can improve outcomes in human HF

## Acknowledgements

Graphical figure was created with BioRender.com.

## Author contributions

ZG contributed to writing the manuscript, conception, design and performed experiments. CVR, AP, DRR, MO, EC, AG, TR, HE, AD, AK, KH, SVS and SH performed experiments and assisted with writing the manuscript. XM provided technical assistance and critically reviewed the manuscript. AA, MSC, KBM and TPC provided human data, and critically reviewed the manuscript. CC performed experiments and critically reviewed the manuscript. JSP and AD provided critical reagents and critically reviewed the manuscript. JAC, LAC, LZ, DG, FRV, AAD, LNP, CB, NOS, MPR and JFD critically reviewed the manuscript. AJ designed experiments and wrote the manuscript.

## Non-standard Abbreviations and Acronyms

ApoM: apolipoprotein M
ApoA-I: apolipoprotein A-I
HDL: high-density lipoprotein
HF: heart failure
LAMP1: lysosomal associated membrane protein 1
LVEF: left ventricular ejection fraction
S1P: sphingosine-1-phosphate
S1PR3: S1P receptor 3
TFEB: transcription factor EB

---

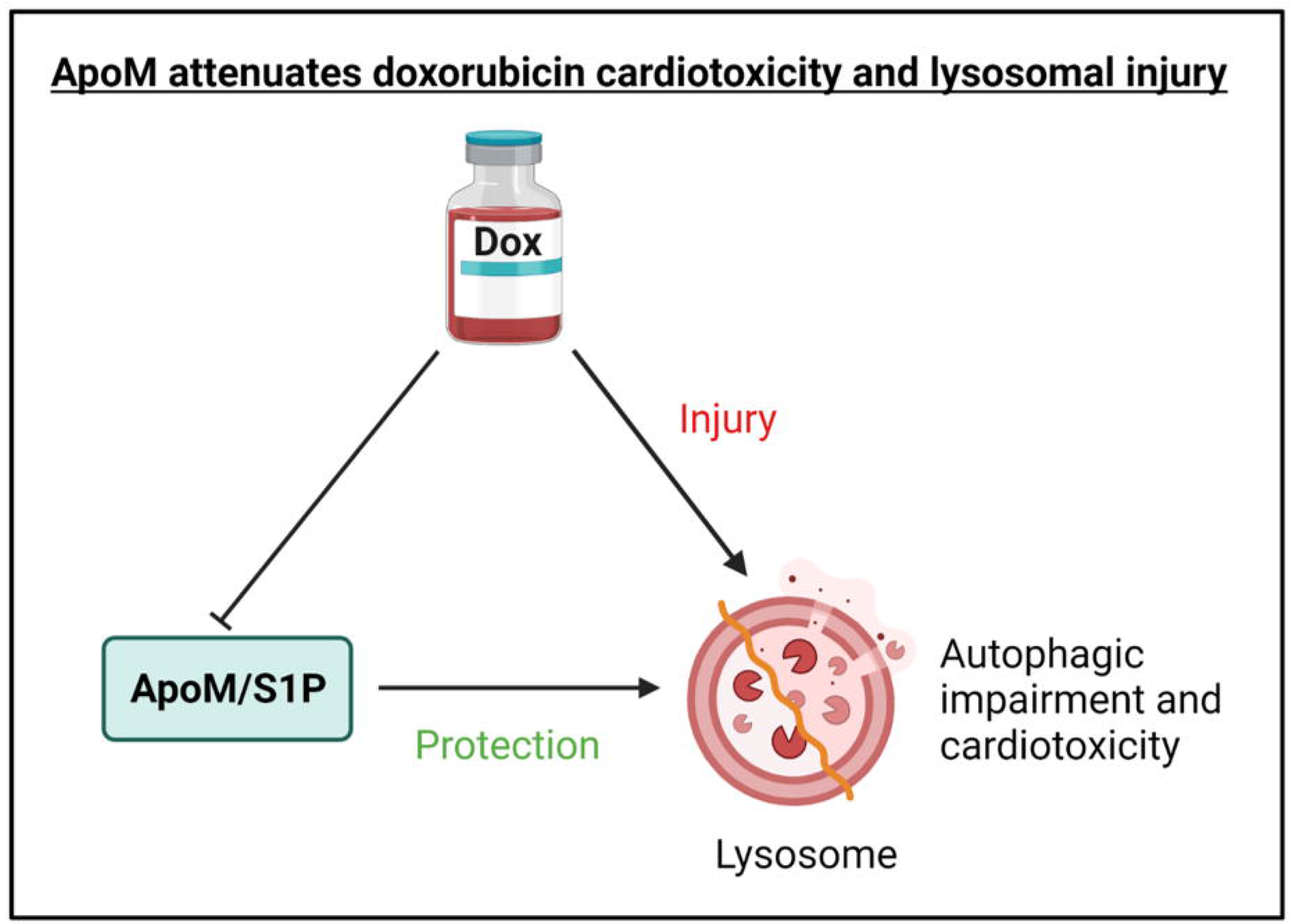

