## Supplementary for "Apolipoprotein M attenuates anthracycline cardiotoxicity and lysosomal injury"

### SUPPLEMENTAL MATERIAL

#### Supplementary methods

##### *Rodent studies*

The Rosa-LAMP1-RFP lysosomal reporter mouse line was generated by a combination of TALEN targeting of the ROSA26 locus followed by homologous recombination with a repair template containing the LAMP1-RFP construct. We first generated a DNA construct consisting of the coding sequence of the rat LAMP1 cDNA connected in frame with the sequence for red fluorescent protein (RFP), followed by sequences for two FLAG epitope tags, a TEV cleavage sequence, a HA epitope tag sequence, and a stop codon sequence. We then inserted this transgene downstream of a CAG promoter and a transcriptional stop cassette flanked by LoxP sites. The CAG-loxP-stop-loxP-LAMP1-RFP construct was then cloned into a targeting vector where it was flanked by homology arms specific to the intronic sequences between exons 1 and 2 of the ROSA26 locus. TALENs specific to this intron were then designed using the Zifit targeter (http://zifit.partners.org). TALEN sequences were assembled using the REAL Assembly TALEN Kit (Addgene 1000000017) and cloned into pJDS71. mRNA for TALENs were generated using these plasmids by in vitro transcription using the mMachine T7 Ultra kit (Ambion). TALEN mRNAs and the final linearized targeting vector were then injected into the pronucleus of C57Bl6/JxCBA hybrid embryos. Founders were screened by PCR to identify mice with insertions into the ROSA26 locus. Founder mice with insertions were bred to C57BL/6J mice to generate F1 offspring which were screened again to confirm germline transmission. F1 mice were subsequently backcrossed with C57BL/6J mice; Rosa-LAMP1-RFP mice were 5 generations into C57BL/6J background in the studies detailed here. Targeting vector construction and TALEN generation were done by the Hope Center Transgenic Vectors Core at Washington University; pronuclear injection was performed by the Mouse Genetics Core at Washington University.

ApoM-CTR and ApoM-TM mice were generated as a knock-in model using human cDNA coding for wild type human ApoM or human ApoM with amino acid changes at F71W, R98A, and R116A respectively. The donor plasmid was synthesized by Eurofins and introduced into C57B6 mouse embryonic stem cells by the Crispr/Cas9 system in the Transgenic Core Facility at University of Copenhagen. Founders were screened by PCR and Southern Blot.

***Blood collection***

Blood was obtained from mice by puncturing the right mandibular vein with a 5.5-mm animal lancet and collecting the blood directly into plasma separator tubes kept on ice and centrifuged at 10,000 g for 5 min. Plasma was transferred to cryovials and snap frozen in liquid nitrogen and stored at -80°C.

***Flow cytometry***

Erythrocytes were removed from blood samples using ammonium chloride-potassium bicarbonate lysis and WBCs were re-suspended in staining buffer (PBS supplemented with 0.5% bovine serum albumin and 2 mM EDTA). Cells were incubated for 30 min at room temperature with pre-titrated saturating dilutions of the following fluorochrome-labeled monoclonal antibodies (BD Biosciences; clone designated in parenthesis): CD45 (104), CD34 (RAM34), CD117 (2B8), and Ly6C (HK1.4). Dead cells were excluded by staining with 2 µg/mL 7-amino-actinomycin D (BD Biosciences) for 5 min prior to analysis. Samples were analyzed on a Gallios flow cytometer (Beckman Coulter), and data were analyzed using FlowJo software (TreeStar, Ashland, OR, USA). APL blasts co-expressed CD34, CD117, and Ly6C.

***Cell culture***

The atria and great vessels were trimmed off, and tissue was finely minced, followed by sequential digestion with 0.5 mg/mL collagenase (Worthington Biochemical, Lakewood, NJ, USA). Ventricular cardiomyocytes were separated from fibroblasts by differential plating and cultured in gelatin-coated tissue culture plates (different plates used depending on the experimental requirements) in medium containing modified Gibco™ Dulbecco's Modified Eagle Medium

(DMEM) (Gibco, 11965-084 ), 10% horse serum (Thermo Fisher, 16050-122), 5% Fetal Bovine Serum (FBS) (Sigma-Aldrich, F2442, St. Louis, MO, USA), 100 µM bromodeoxyuridine (Brdu), 100 U/mL penicillin, 100 µg/mL streptomycin (Thermo Fisher, 15140148) and L-glutamine (Sigma-Aldrich, G8540). Neonatal rat cardiomyocytes were serum-starved overnight, then treated with vehicle or FTY720 (Cayman Chemicals, 10006292, Ann Arbor, MI, USA) prior to treating with 0.5 mM doxorubicin for 4 hr. Cells were incubated with 1 µM Lysosensor Yelow/Blue DND 160 (Thermo Fisher, L7545) for 3 min, washed with HCSS and imaged by collecting pairs of images excited at 380 nm and 340 nm (Polychrome V, FEI, Munjch, Germany) through a long pass 480 nm emission filter through a 40x/1.35 oil immersion lens (Olympus) using an iMIC microscope (FEI). After subtracting the matching wavelength background, the images were divided by each other to yield an excitation ratio images. The ratio values of individual lysosomes were determined using Live Acquisition software package (FEI) and converted to pH using a calibration curve prepared by measuring Lysosensor excitation ratio on calibration buffers of known pH (3.0-6.5) on the same optical system.

***Quantitative Real-Time Polymerase Chain Reaction analysis (qPCR)***

Total RNA in cardiac tissues were extracted using RNeasy Mini kit (Qiagen, #74104, Hilden, Germany), and the first-strand cDNA was prepared using the iScriptTM cDNA synthesis kit (Bio-Rad, #1708890, Hercules, CA, USA). qPCR analysis was performed with SYBR Green Master Mix (Bio-Rad, #1725121) on QuantStudio 3 Real-Time PCR system (Applied Biosystems, A28136, Foster City, CA, USA) to examine the relative mRNA levels of indicated genes. Mice sequences for qRT-PCR primers are shown below: ***Tfeb***: forward 5'-GTC TAG CAG CCA CCT GAA CGT-3', reverse 5'-ACC ATG GAG GCT GTG ACC TG-3'; ***Actb***: forward 5'-CAG AAG GAG ATC ACT GCC CT-3', reverse 5'-AGT ACT TGC GCT CAG GAG GA-3'; ***36b4***: forward 5'-GCT TCG TGT TCA CCA AGG AGG A-3', reverse 5'-GTC CTA GAC CAG TGT TCT GAG C-3'; ***Rpl32***: forward 5'-CCT CTG GTG AAG CCC AAG ATC-3', reverse 5'-TCT GGG TTT CCG CCA GTT T-

3'; ***Gapdh***: forward 5'-ACT CCC ACT CTT CCA CCT TC-3', reverse 5'-TCT TGC TCA GTG TCC TTG C-3'.

***Western blot analysis***

Myocardial extracts were prepared by homogenization of ventricular tissue with lysis buffer (composition in mM: 50 Tris HCl, pH 7.4; 2.5 EDTA; 10 EGTA; 20 NaF; 25 $Na_4P_2O_7.10\ H_2O$; 2 $Na_3VO_4$; 25 NaCl) containing 0.2% NP-40, 1x PI and 1x PPI. 15-40 µg of protein were separated by sodium dodecyl sulfate polyacrylamide gel electrophoresis (SDS-PAGE) (Bio-Rad) and transferred to a polyvinylidene fluoride (PVDF) membrane (Millipore, #3010040001). After being blocked with 5% fat-free milk in TBST, the membranes were incubated with individual primary antibodies overnight at 4°C. Subsequently, the membrane was incubated with a horseradish peroxidase-conjugated secondary antibody (Cell Signaling, #7074 or #7076), and exposed to ClarityTM Western ECL Substrate (Bio-Rad, #170-5060) using the ChemiDoc System (Bio-Rad, 12003153) for detection of protein expression. Primary antibodies employed were as follows: **mouse ApoM** (LS Bio, C158166, diluted 1:1000, Seattle, WA, USA), **human ApoM** (R&D, AF4550, diluted 1:1000, Minneapolis, MN, USA), **TFEB** (Bethyl Labs, A303-673A, diluted 1:1000, Montgomery, TX, USA), **AKT** (Cell Signaling, #9272s, diluted 1:1000), **phospho-AKT(Ser473)** (Cell Signaling, #4060s, diluted 1:2000), **Histone H3** (Cell Signaling, #9715s, diluted 1:1000), **GAPDH** (Abcam, ab22555, diluted 1:5000, Cambridge, MA, USA), **Albumin** (Cell Signaling, #4929s, diluted 1:1000) and **β-actin** (Sigma-Aldrich, A2066, diluted 1:4000). The band intensity was measured and analyzed with ImageJ software (Bio-Rad).

***Cytoplasm and Nuclear protein extraction***

Heart tissue were homogenized in 1x isotonic lysis buffer (ILB) containing 10 mM dithiothreitol (DTT), 1x Protease Inhibitor (PI) (Millipore, #4693132001, Burlington, MA, USA) and 1x Halt Protease and Phosphatase Inhibitor (PPI) (Thermo Fisher, #78446) using the Mechanical Homogenizers (IKA, #3737001, Staufen, Germany) and centrifuged at 4°C, 10,500 g for 15 min, and the supernatants were cytoplasmic fraction. The pellets were washed 3 times with 1x ILB and fully resuspended in 1x extraction buffer (EB) containing 10 mM DTT, 1x PI and 1x PPI. The suspension was kept on the ice for 30 min and then completely sonicated by the Sonifier (Branson Ultrasonics, #250-450, Danbury, CT, USA). The supernatant from the 20,500 g spin at 4°C for 5 min were the nuclear fraction. Expression of proteins localized to the nucleus (Histone H3) and cytoplasm (GAPDH) was examined to confirm relative enrichment.

***Insoluble protein isolation***

Heart tissue was mechanically homogenized with a Dounce homogenizer in 500-1000 uL of homogenization buffer (0.3 M KCl, 0.1 M $KH_2PO_4$, 50 mM $K_2HPO_4$, 10 mM EDTA, 4 mM Na Orthovanadate, 100 mM NaF, Protease inhibitor, pH to 6.5). Homogenized samples were passed through mesh basket on ice, followed by collection of the lysate run-through which was incubated on ice for 30 min. A known volume of the sample was transferred to another Eppendorf tube and 20% NP-40 was added to for a final concentration of 1% NP-40. Samples were then incubated on ice for 30 min, and spun at 20,500 g for 15 min, 4°C. Supernatant was collected as soluble fraction. The pellet was washed 3 times with cold PBS (following addition of 1 mL PBS to each pellet, and spin down at 20,500 g for 10 min) followed by resuspension in 1% SDS, 10 mM Tris buffer to generate the insoluble fraction.

**Supplementary Tables**

**Supplementary Table 1**

**General characteristics of PHFS study participants with anthracycline-induced cardiomyopathy (CMP) vs non-ischemic cardiomyopathy.**

| Clinical variable | Anthracycline CMP (n=46) | Non-ischemic CMP (n=1447) | p value |
|---|---|---|---|
| ApoM (AU) | 862 (738, 999) | 840 (701, 996) | 0.8119 |
| Age (years) | 56.1 (43.5, 62.6) | 55 (44, 64) | 0.7149 |
| Male sex | 18 (39.13%) | 864 (59.71%) | 0.0052 |
| Race/Ethnicity | | | |
| Caucasian | 33 (71.74%) | 983 (67.93%) | 0.5858 |
| African American | 10 (21.74%) | 388 (26.81%) | 0.4435 |
| Asian | 0 (0.00%) | 15 (1.04%) | 1.0000 |
| BMI (kg/m$^2$) | 27.4 (24.6, 30.3) | 28.9 (25.1, 34.5) | 0.0661 |
| Systolic BP (mmHg) | 110 (100, 124) | 114 (100, 130) | 0.1361 |
| Diastolic BP (mmHg) | 70 (60, 77.5) | 70 (62, 78) | 0.5646 |
| Current smoking | 1 (2.17%) | 138 (9.54%) | 0.1183 |
| Diabetes | 7 (15.22%) | 333 (23.01%) | 0.2145 |
| Atrial fibrillation or flutter | 11 (23.91%) | 470 (32.48%) | 0.2209 |
| LV ejection fraction (%) | 36.3 (25, 52.5) | 35 (20, 50) | 0.2640 |
| History of stent | 4 (8.70%) | 89 (6.15%) | 0.5267 |
| History of CABG | 1 (2.17%) | 58 (4.01%) | 1.0000 |
| eGFR | 65.3 (50.3, 72.7) | 59.6 (45.4, 73.5) | 0.4564 |
| NYHA Class | | | |
| NYHA 1 | 9 (19.57%) | 295 (20.54%) | 0.8715 |
| NYHA 2 | 23 (50.00%) | 662 (46.10%) | 0.6015 |
| NYHA 3-4 | 14 (30.43%) | 479 (33.36%) | 0.6789 |
| Medication Use | | | |
| Beta Blocker | 37 (80.43%) | 1274 (88.04%) | 0.1204 |
| Aspirin | 23 (50.00%) | 658 (45.47%) | 0.5440 |
| ACEI/ARBs | 36 (78.26%) | 1237 (85.49%) | 0.1734 |
| Hydralazine | 1 (2.17%) | 132 (9.12%) | 0.1181 |
| Nitrates | 1 (2.17%) | 169 (11.68%) | 0.0457 |
| Statin | 13 (28.26%) | 575 (39.74%) | 0.1168 |
| CCBs | 1 (2.17%) | 148 (10.23%) | 0.0797 |
| Warfarin | 9 (19.57%) | 493 (34.07%) | 0.0404 |
| Insulin | 1 (2.17%) | 148 (10.23%) | 0.0797 |

**Supplementary Table 2**

***Clinical characteristics of breast cancer cardiotoxicity cohort.***

| Clinical variable | Mean ± SE or % |
|---|---|
| Age | 49.3 ± 10.9 |
| Body mass index | 25.5 ± 4.2 |
| Hypertension | 38.9% |
| Hyperlipidemia | 27.8% |
| Diabetes | 2.8% |
| Smoking history | 8.3% |

**Supplementary Figures**

**Supplementary Figure 1**

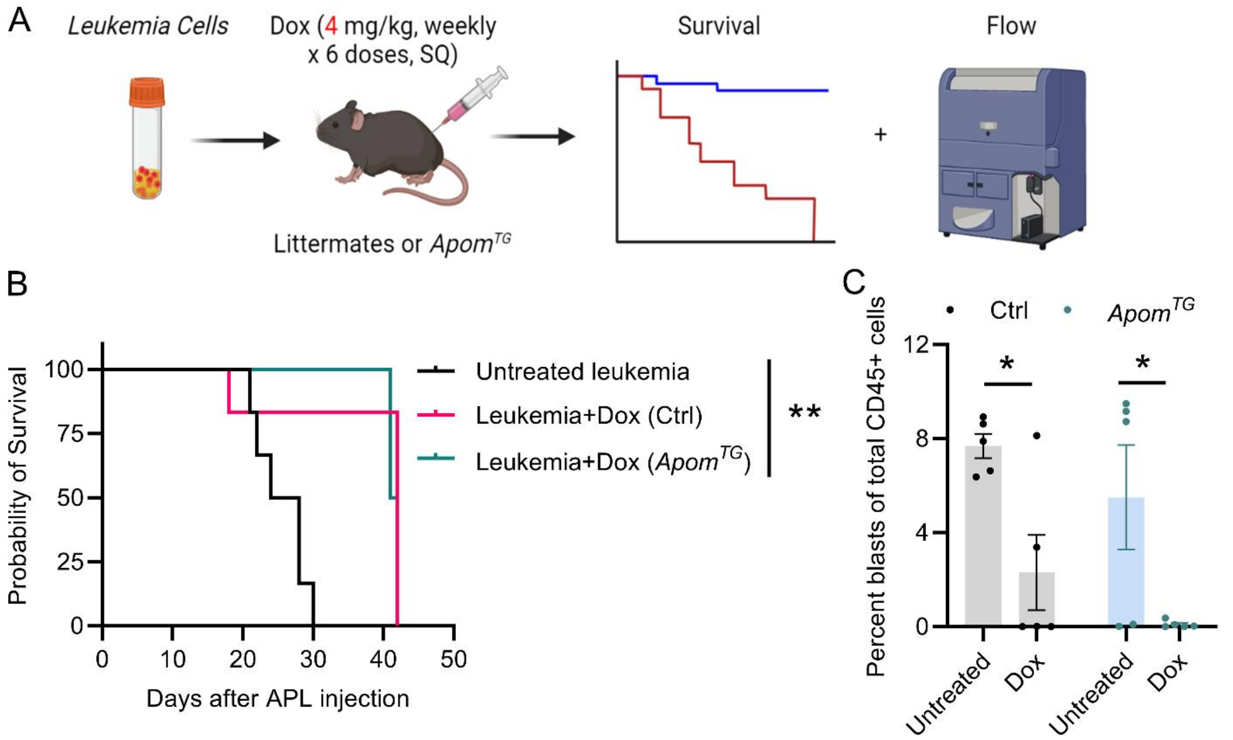


**Supplementary Figure 1. ApoM does not attenuate the anti-leukemic efficacy of doxorubicin in vivo.**

**A)** Schematic diagram of the experimental. **B)** Survival of littermate control (Ctrl) and *Apom*$^{TG}$ mice injected with APL cells and treated with vehicle (black untreated control) or doxorubicin (blue and red, 4 mg/kg doxorubicin (Dox) SQ x 6 doses). Log-rank test, n=6 untreated, 6 Ctrl, and 4 *Apom*$^{TG}$. **C)** Percentage of APL cells in the peripheral blood was measured by flow cytometry after 6 doses of Dox in Ctrl and *Apom*$^{TG}$ mice on day 17 after APL injection. Two-way ANOVA with Sidak's correction for multiple comparisons, n=5 per group. Each dot represents one mouse. *$p < 0.05$, **$p < 0.01$.

**Supplementary Figure 2**

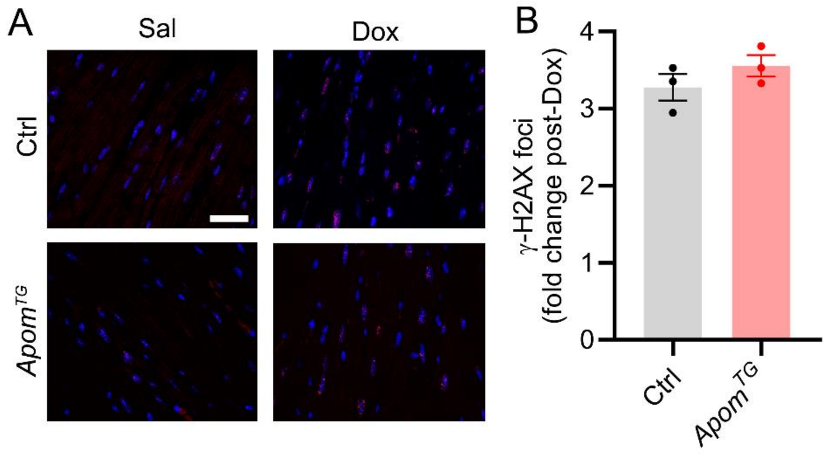


**Supplementary Figure 2. Anti-γ-H2AX staining in littermate control (Ctrl) and *Apom*$^{TG}$ mice 48 hr after treatment with 10 mg/kg doxorubicin (Dox) IP.**

**A)** Representative images of myocardial sections were stained with anti-γ-H2AX (red) and DAPI (blue), scale bar=20 μm. **B)** Blinded quantification of (A). Student's t-test, n=3 per group. Each dot represents one mouse.

**Supplementary Figure 3**

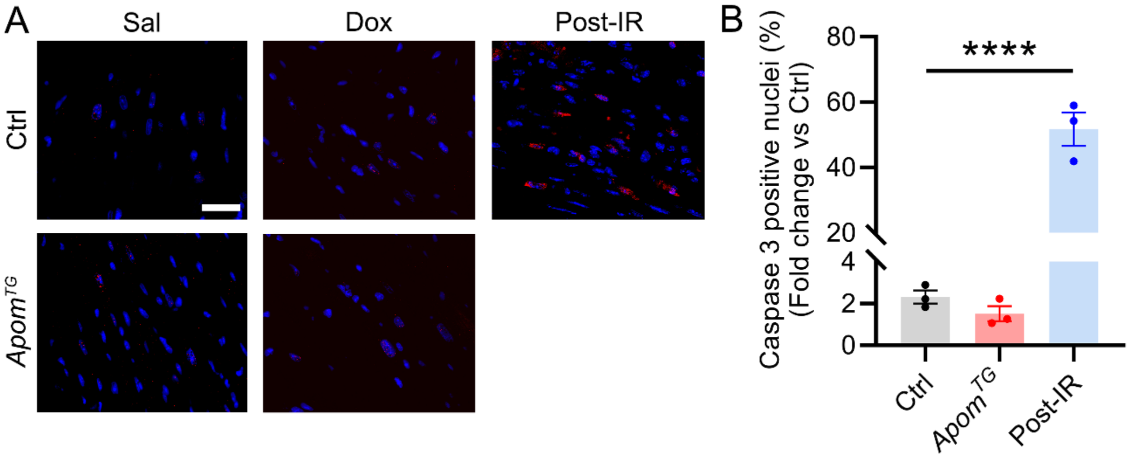


**Supplementary Figure 3. Anti-caspase 3 staining in littermate control (Ctrl) and *Apom$^{TG}$* mice 48 hr after treatment with 10 mg/kg doxorubicin (Dox) IP.**

**A)** Representative images of myocardial sections were stained with anti-caspase 3 (red) and DAPI (blue), scale bar=20 µm. **B)** Blinded quantification of (A). One-way ANOVA with Dunn's correction for multiple comparisons, n=3 per group. Post-ischemia reperfusion (IR) sections were utilized from mice 28 days post-90 min closed-chest IR of the left anterior descending artery as a positive control. Each dot represents one mouse. ****$p < 0.0001$.

**Supplementary Figure 4**

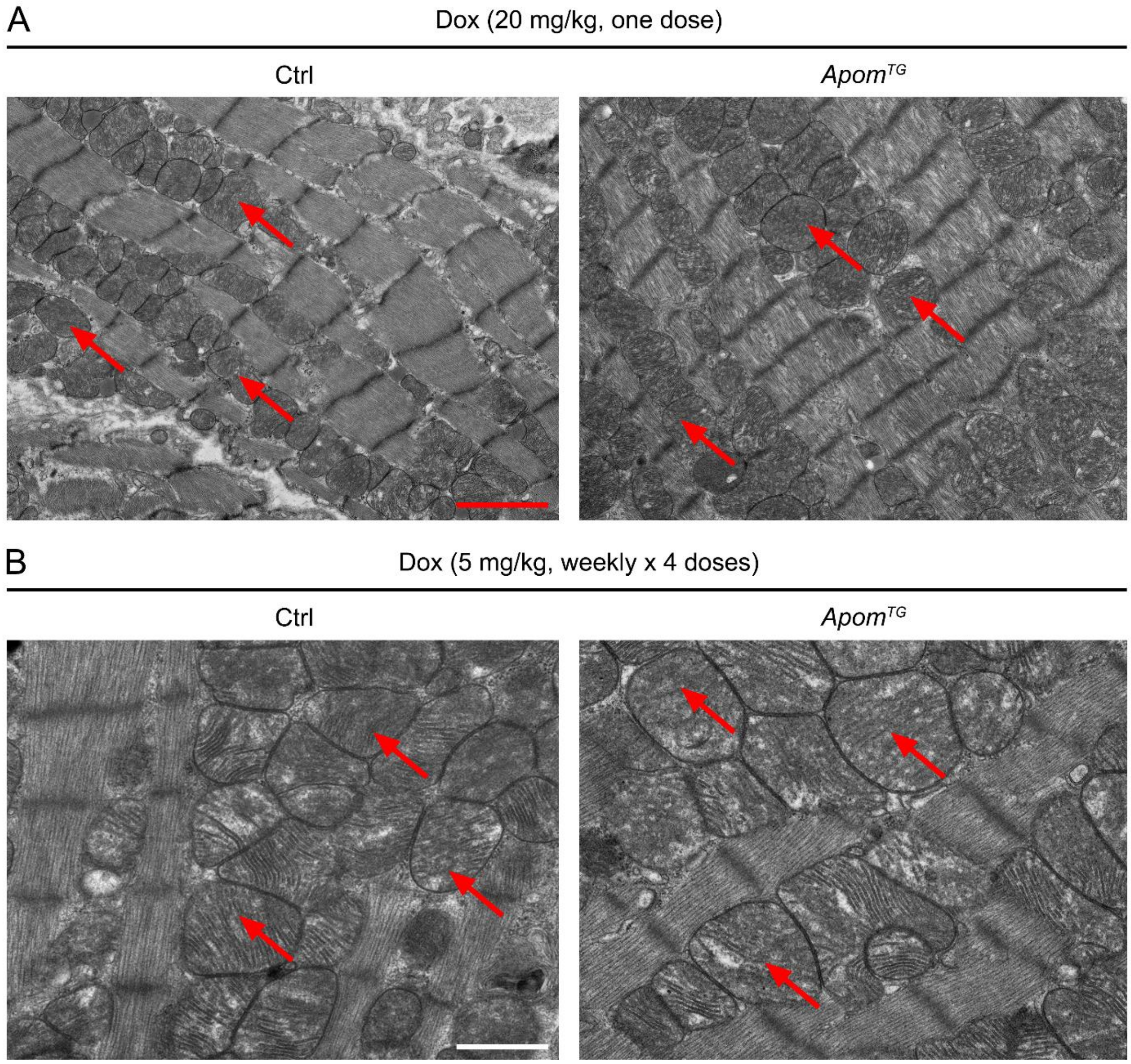


**Supplementary Figure 4. ApoM does not improve mitochondrial ultrastructure in doxorubicin cardiotoxicity.**

**A)** Representative images of transmission electron micrograph of myocardium from littermate control (Ctrl) and *Apom*$^{TG}$ mice treated with 20 mg/kg doxorubicin for 5 days, red scale bar=2 μm. **B)** Representative images of transmission electron micrograph of myocardium from Ctrl and *Apom*$^{TG}$ mice 3 months after receiving 5 mg/kg doxorubicin once weekly x 4 doses, IV, white scale bar=50 nm. Red arrows indicate rarified mitochondrial cristae.

**Supplementary Figure 5**

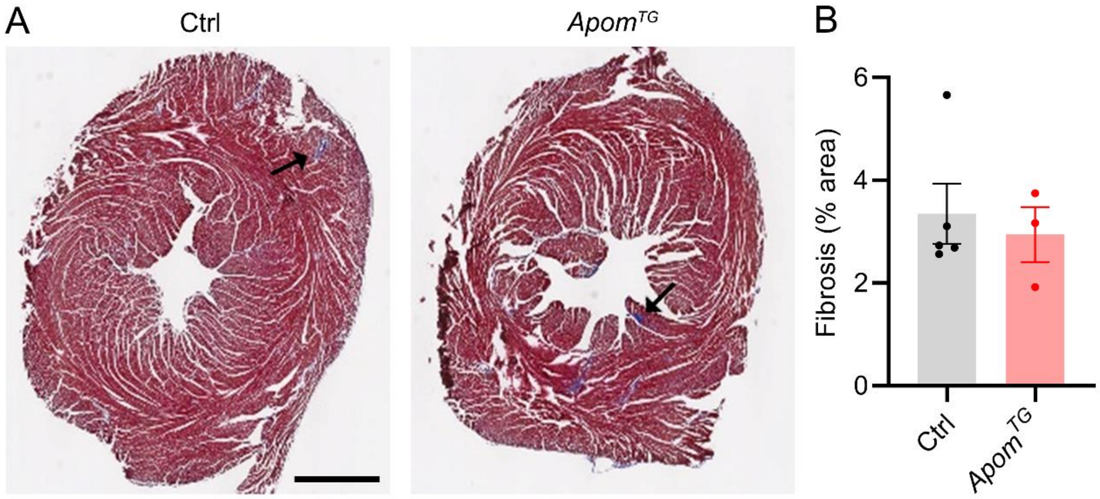


**Supplementary Figure 5. ApoM does not prevent myocardial fibrosis in doxorubicin cardiotoxicity.**

**A)** Representative images of Masson's trichrome staining of myocardial sections from littermate control (Ctrl) and *Apom*$^{TG}$ mice 3 months after treatment with 5 mg/kg doxorubicin weekly x 4 doses, scale bar=1 mm. Black arrows indicate fibrosis. **B)** Blinded quantification of (A). Student's t-test, n=5 Ctrl vs n=3 *Apom*$^{TG}$. Each dot represents one mouse.

**Supplementary Figure 6**

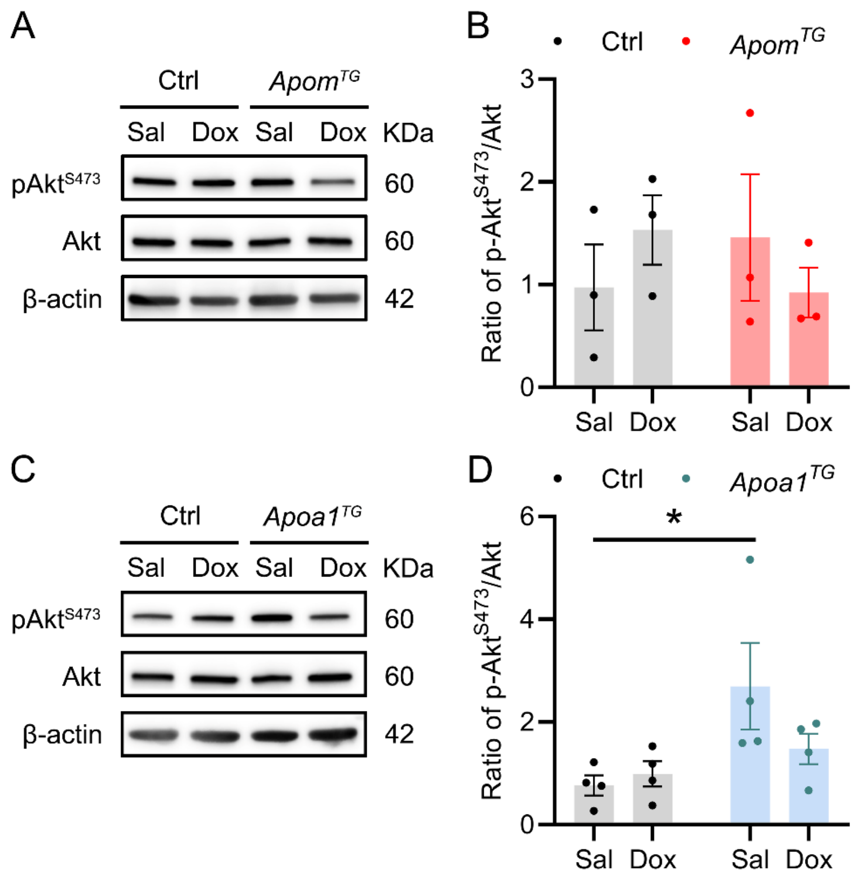


**Supplementary Figure 6. *Apom$^{TG}$* mice do not exhibit increased myocardial Akt phosphorylation.**

**A)** Representative western blot for myocardial phospho-Akt (Ser 473) and total Akt from littermate control (Ctrl) and *Apom$^{TG}$* mice 48 hr after treatment with saline (Sal) or 10 mg/kg doxorubicin (Dox) IP. **B)** Quantification of ratio of p-AktS473/Akt from (A). Two-way ANOVA with Sidak's correction for multiple comparisons, n=3 per group. **C)** Representative western blot for myocardial phospho-Akt (Ser 473) and total Akt from control and *Apoa1$^{TG}$* mice 48 hr after treatment with vehicle or 10 mg/kg doxorubicin IP. **D)** Quantification of ratio of p-AktS473/Akt from (C). Two-way ANOVA with Sidak's correction for multiple comparisons, n=4 per group. Each dot represents one mouse. *$p < 0.05$.

**Supplementary Figure 7**

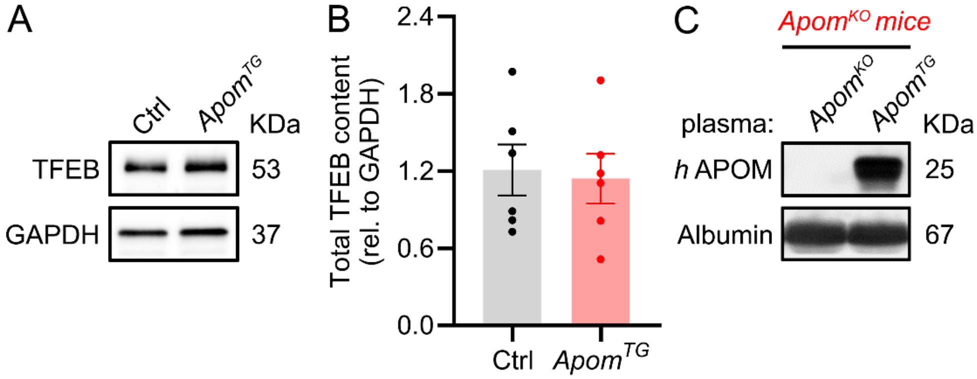


**Supplementary Figure 7. ApoM does not affect the total protein level of TFEB in the myocardium.**

**A)** Representative western blot of TFEB from isolated myocardial total protein extracts from littermate control (Ctrl) and *Apom*$^{TG}$ mice. **B)** Quantification of total TFEB from (A). Student's t-test, n=6 per group. Each dot represents one mouse. **C)** Representative western blot of human ApoM from plasma obtained from *Apom*$^{KO}$ mice 2 days after transfer of 120 μL plasma obtained from *Apom*$^{KO}$ or *Apom*$^{TG}$ donor mice.

**Supplementary Figure 8**

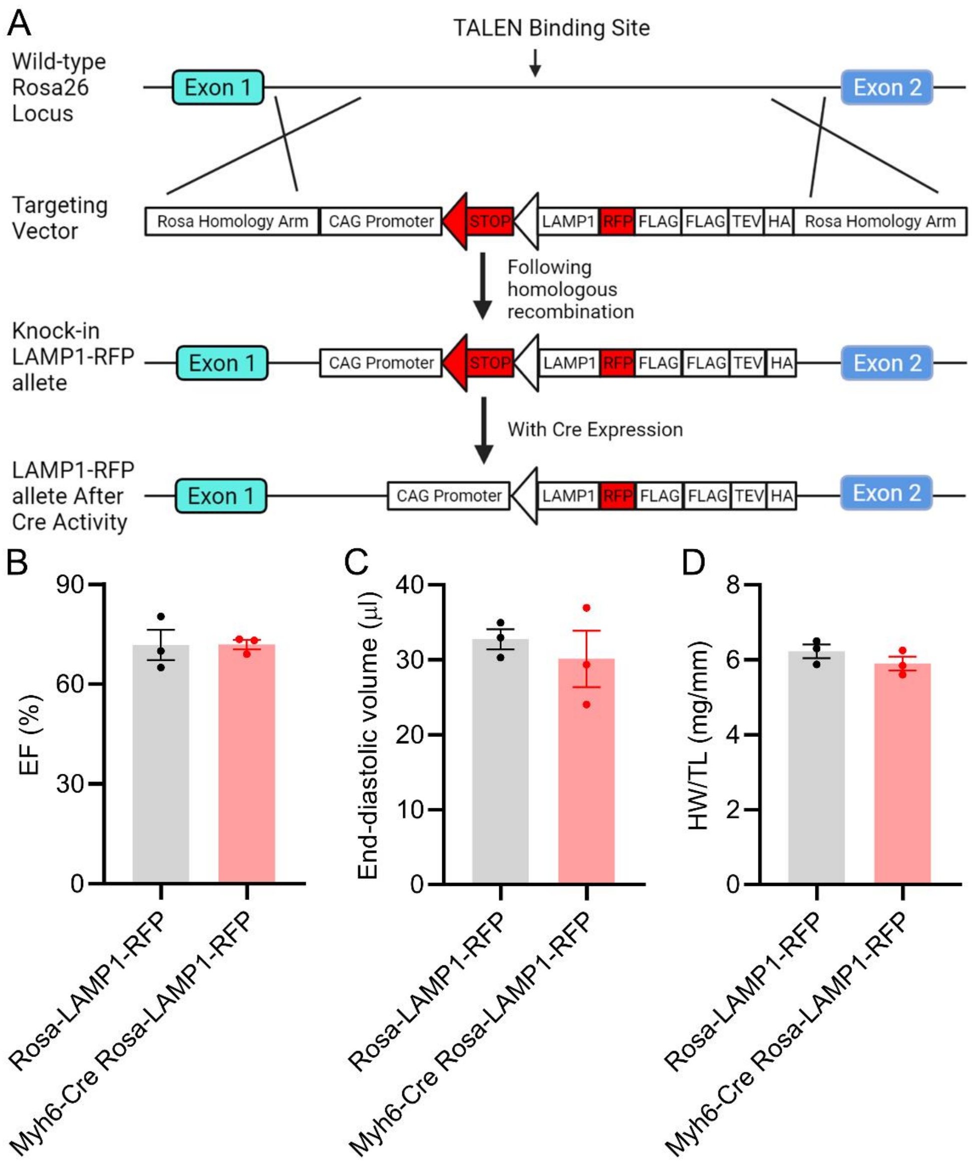


**Supplementary Figure 8. Echocardiographic and morphometric parameters in Rosa-LAMP1-RFP lysosomal reporter allele mice at 8 weeks old.**

**A)** Schematic of targeting strategy for generating the Rosa-LAMP1-RFP lysosomal reporter allele. **B-C)** Assessment of ejection fraction (B) and left ventricular volume (C) by echocardiography of *Myh6-Cre* Rosa-LAMP1-RFP hearts versus controls. Student's t-test, n=3 per group. Each dot represents one mouse. **D)** Heart weight in *Myh6-Cre* Rosa-LAMP1-RFP hearts versus control hearts, normalized to tibia length. Student's t-test, n=3 per group. Each dot represents one mouse.
